# Medial Prefrontal Cortex Serotonin Input Regulates Cognitive Flexibility in Mice

**DOI:** 10.1101/2023.03.30.534775

**Authors:** Ashlea A. Morgan, Nuno D. Alves, Gregory S. Stevens, Tamanna T. Yeasmin, Alexandra Mackay, Saige Power, Derya Sargin, Carla Hanna, Arwa L. Adib, Annette Ziolkowski-Blake, Evelyn K. Lambe, Mark S. Ansorge

## Abstract

The medial prefrontal cortex (mPFC) regulates cognitive flexibility and emotional behavior. Neurons that release serotonin project to the mPFC, and serotonergic drugs influence emotion and cognition. Yet, the specific roles of endogenous serotonin release in the mPFC on neurophysiology and behavior are unknown. We show that axonal serotonin release in the mPFC directly inhibits the major mPFC output neurons. In serotonergic neurons projecting from the dorsal raphe to the mPFC, we find endogenous activity signatures pre-reward retrieval and at reward retrieval during a cognitive flexibility task. *In vivo* optogenetic activation of this pathway during pre-reward retrieval selectively improved extradimensional rule shift performance while inhibition impaired it, demonstrating sufficiency and necessity for mPFC serotonin release in cognitive flexibility. Locomotor activity and anxiety-like behavior were not affected by either optogenetic manipulation. Collectively, our data reveal a powerful and specific modulatory role of endogenous serotonin release from dorsal raphe-to-mPFC projecting neurons in cognitive flexibility.

## Introduction

Organisms adapt to survive. Both cognitive flexibility and emotionality have evolved to facilitate adaptation and both domains are dependent on medial prefrontal cortex (mPFC) function.

Cognitive flexibility facilitates adaptation by allowing us to change cognitive patterns in response to changing external stimuli (Dennis et al., 2010). Humans with lesions to the mPFC exhibit impairments in aspects of cognitive flexibility requiring shifts in attention (Shallice et al., 2007, Stuss, 2011). In marmoset monkeys, neurotoxic lesions to the mPFC worsen performance in a cognitive flexibility task where attention must shift to a previously irrelevant dimension (extradimensional set shift) (Heilbronner et al., 2016). Pharmacological inactivation or lesions of the rodent mPFC impair performance in tasks that assess cognitive flexibility (Bissonette et al., 2008, Bissonette et al., 2013, Brigman et al., 2013, Cho et al., 2015, Darrah et al., 2008, Marquardt et al., 2014, Roy et al., 2010, Troudet et al., 2016). Likewise, optogenetic silencing of mPFC pyramidal neurons through activation of interneurons impairs cognitive flexibility in an attention set shifting task (Biro et al., 2019, Marton et al., 2018). In mice, prefrontal cortex neurons encode recent trial outcomes and enable attentional set-shifting by relaying feedback information to downstream targets (Spellman et al., 2021).

Emotions are internal experiences triggered by external events or cognitive processes, leading us to seek or avoid places, situations, or circumstances. The human mPFC is tightly associated with emotional regulation (Anderson et al., 2006, Barrash et al., 2000, Bechara et al., 1994, Damasio et al., 1990, Kroes et al., 2019, Szczepanski et al., 2014). Likewise in mice, the mPFC is required for responses in anxiogenic environments, with pharmacological and physical lesions reducing anxiety-sensitive behaviors (Deacon et al., 2003, Klein et al., 2010, Lacroix et al., 2000, Lisboa et al., 2010, Rebello et al., 2014, Shah et al., 2003, Sullivan et al., 2002). Furthermore, deep layer neurons in the prelimbic sub-region (PL) of the mPFC create task-related safe and aversive representations in an anxiogenic environment (Adhikari et al., 2011).

The mPFC receives dense serotonergic input, which acts as a powerful modulator of neuronal activity (Celada et al., 2013, Chandler et al., 2013, Sargin et al., 2019). Exogenous serotonin or related agonist administration inhibits mPFC pyramidal cells and glutamatergic input into the mPFC, and activates mPFC fast-spiking interneurons (Elliott et al., 2018, Kjaerby et al., 2016, Tian et al., 2016, Zhong et al., 2011). However, which neurophysiological consequences are triggered upon endogenous serotonin release is unknown.

Optogenetic photostimulation of DR-mPFC serotonergic neurons promotes waiting when the timing of reward delivery is uncertain (Miyazaki et al., 2020). Furthermore, 5-HT_1A_ receptor agonist infusion into the PL improves accuracy on an attention task in rodents (Winstanley et al., 2003), and serotonin depletion in the PFC of marmosets impairs cognitive flexibility (Clarke et al., 2004, Clarke et al., 2005). However, the consequences of endogenous serotonin release on cognitive flexibility are not known (Dias et al., 1996a, Dias et al., 1996b, Rogers et al., 2000).

Lastly, drugs that target the serotonin system are typically prescribed to treat disorders of emotional dysregulation (Blier et al., 1990, Nutt, 2005). Exposure to anxiogenic environments increases extracellular serotonin as measured via microdialysis in the PFC of guinea pigs (Rex et al., 1998), and 5-HT_1A_ receptor agonist infusion into the PL induces anxiolytic effects in rats (Yamashita et al., 2018). These findings indicate that cortical serotonin release reduces anxiety-like behavior. Yet again, it is unknown how serotonergic neurons projecting to the mPFC respond to threat and if the endogenous terminal release of serotonin in the mPFC modulates anxiety.

To address these unknowns, we investigated the role of endogenous serotonin input into the mPFC in modulating neuronal activity and behavior in mice. Specifically, after first determining the origin of mPFC serotonin input using viral retrograde tracing, we assessed how terminal release of serotonin affects mPFC pyramidal neuron activity using whole-cell electrophysiology in acute mouse brain slices. Through *in vivo* fiber photometry, we then evaluated the activity of serotonergic neurons projecting to the mPFC during behavior. Lastly, through *in vivo* optogenetics, we assessed the modulatory role of serotonergic input into the mPFC on anxiety-like behavior as well as intradimensional rule reversal and extradimensional rule shift performance in an attention set-shifting task.

## Results

### Evoked PL serotonin terminal release hyperpolarizes layer V (LV) pyramidal neurons

We first sought to identify the source of mPFC serotonergic inputs, using retrograde viral tracing. We injected the Cre-dependent retrograde HSV reporter virus HSV-hEF1α-LS1L-GFP into the mPFC of SERT-Cre mice (**Supplementary Figure 1A-B**), and 4 weeks later detected GFP^+^ 5-HT^+^ cells in the dorsal raphe (DR) (**Supplementary Figure 1C**).

Next, to investigate the electrophysiological consequences of endogenous neurotransmitter release from serotonergic axon terminals in the PL, we used an optogenetic approach paired with whole-cell recordings from acute brain slices. We crossed the serotonin specific ePet1-Cre driver line (Scott et al., 2005) with the Ai32 reporter line (Madisen et al., 2012), which Cre-dependently expresses channelrhodopsin-2-enhanced yellow fluorescent fusion protein (ChR2-eYFP). We recorded electrophysiological responses to optogenetic stimulation of serotonergic terminals in coronal sections of ePet1-Cre^-/+^::Ai32^+/+^ mice containing the PL, patching pyramidal neurons in layer V (LV), the major output layer of the prefrontal cortex (**Figure 1A**). Opto-stimulation (10 ms pulses, 20 Hz, 5 s) elicited outward currents in ePet1-Cre^-/+^::Ai32^+/+^ (opto-serotonin elicited current: unpaired t-test t(47) = 5.1, *P* < 0.0001, **Figure 1B,C**) in contrast to littermate control mice (ePet1-Cre^-/-^::Ai32^+/+^). Opto-stimulation elicited outward currents in the presence of synaptic blockers (bicuculline, 3 µM, CNQX, 20 µM, D-APV, 50 µM; **Figure 1D**) that were not significantly different from those elicited in their absence (unpaired t-test *t*(35) = 0.5, *P* = 0.6) suggesting direct inhibition of the major output neurons of mPFC. To further investigate this direct opto-serotonin inhibition, we interrogate the 5-HT_1A_ receptor, an inhibitory receptor broadly expressed and active in the output neurons of the mPFC (Amargos-Bosch et al., 2004, Yin et al., 2017). Application of the selective 5-HT_1A_ receptor antagonist WAY100635 (30 nM, 10 min) significantly inhibited opto-serotonin elicited currents (unpaired t-test *t*(38) = 2.9, *P* = 0.007) in ePet1-Cre^-/+^::Ai32^+/+^mice.

**Figure 1.**
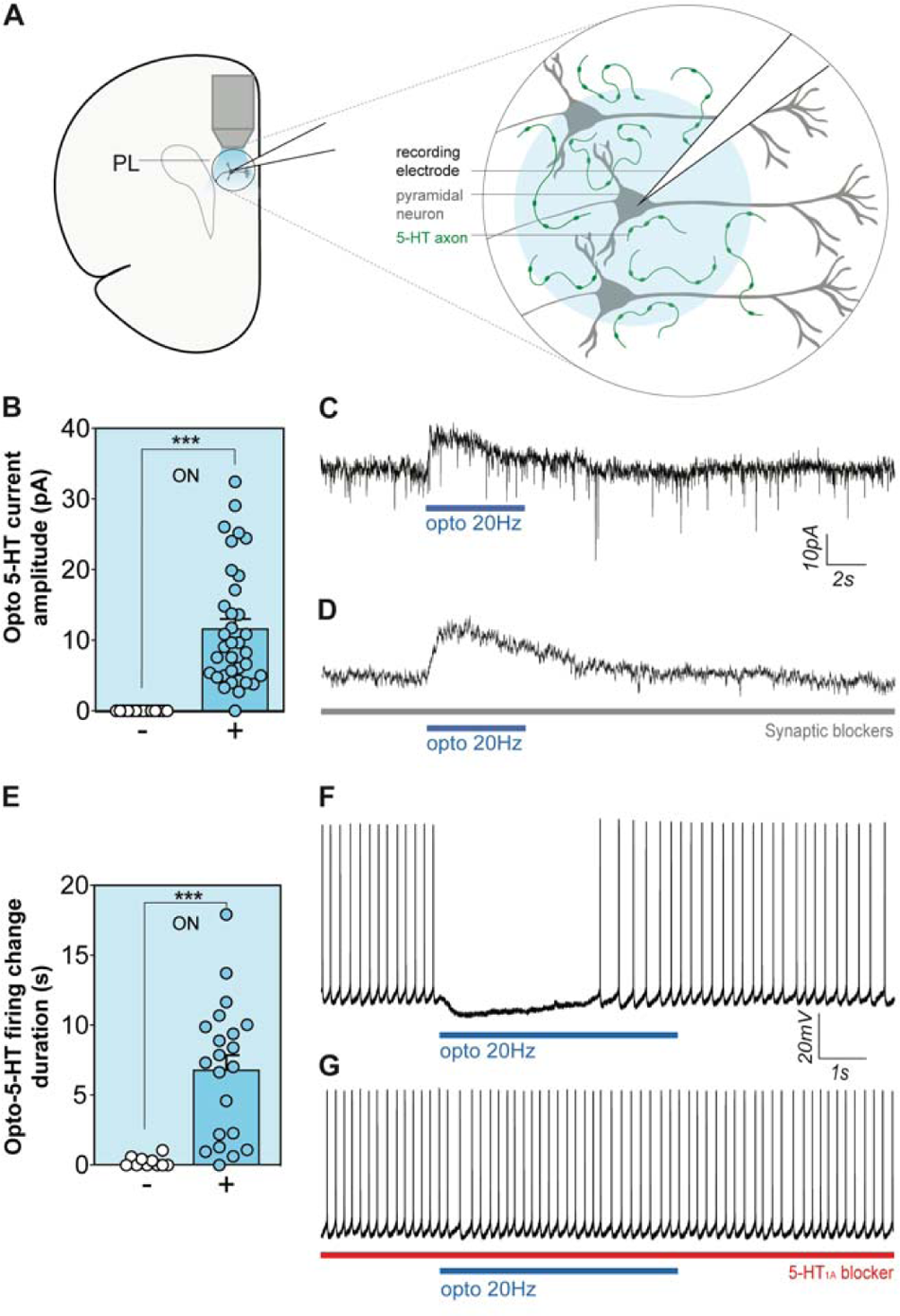
Optogenetic stimulation of serotonergic terminals inhibits the major output neurons of the prelimbic cortex. Opto-stimulation of PL serotonergic terminals (**A**) of ePet1-Cre^-/+^::Ai32^+/+^ mice elicited outward currents in PL pyramidal LV neurons (**B**, **C**) but not Ai32^+/+^ mice, an effect unperturbed by synaptic blockers (**D**). Hyperpolarization of PL pyramidal LV neurons in response to terminal photostimulation suppresses action potential firing (**E**, **F**) in ePet1-Cre^-/+^::Ai32^+/+^ but not ePet1-Cre^-/-^::Ai32^+/+^ mice, an effect blocked by treatment with 5-HT_1A_ receptor antagonist WAY100635 (**G**). Optophysiological recordings were performed in 70 neurons from 15 mice, with Student’s unpaired t-test used to compare groups, \*\*\**P* < 0.001.

To assess the impact of terminal serotonin release on the firing properties of these LV pyramidal neurons, we depolarized patched neurons to elicit action potential firing (3 ± 1 Hz) before applying opto-stimulation. This approach revealed that terminal photostimulation (10 ms pulses, 20 Hz, 5 s) is sufficient to suppress action potential firing of prelimbic (PL) LV pyramidal neurons from ePet1-Cre^-/+^::Ai32^+/+^mice (unpaired t-test *t*(29)= 4.2, *P* = 0.0002, **Figure 1D, E**) but exerts no effect in ePet1-Cre^-/-^::Ai32^+/+^ controls. Consistent with a direct 5-HT_1A_ effect, Application of WAY100635 prevented the optogenetic inhibition of action potential firing of PL LV pyramidal neurons observed in ePet1-Cre^-/+^::Ai32^+/+^ mice (paired t-test t(4) = 8.5, *P* = 0.001; **Figure 1F**).

Together, these results show that direct, 5-HT_1A_-mediated inhibition of the major output neurons of mPFC can be achieved optophysiologically in ePet1-Cre^-/+^::Ai32^+/+^ transgenic mice.

### Specific activity signatures of DR-to-mPFC serotonergic neurons during cognitive flexibility tasks

Next, to investigate the activity of serotonergic neurons projecting to the mPFC during behavior, we expressed the genetically encoded Ca^2+^ sensor *GCaMP6s* using a Cre-dependent retrograde viral approach and recorded bulk fluorescence using fiber photometry. Specifically, we injected HSV-hEF1 α-LSL1L-GCaMP6s into the PL and implanted an optical fiber in the DR. 4 weeks later, mice were recorded in the open-field test and the two-choice digging task (**Figure 2A-D**).

**Figure 2.**
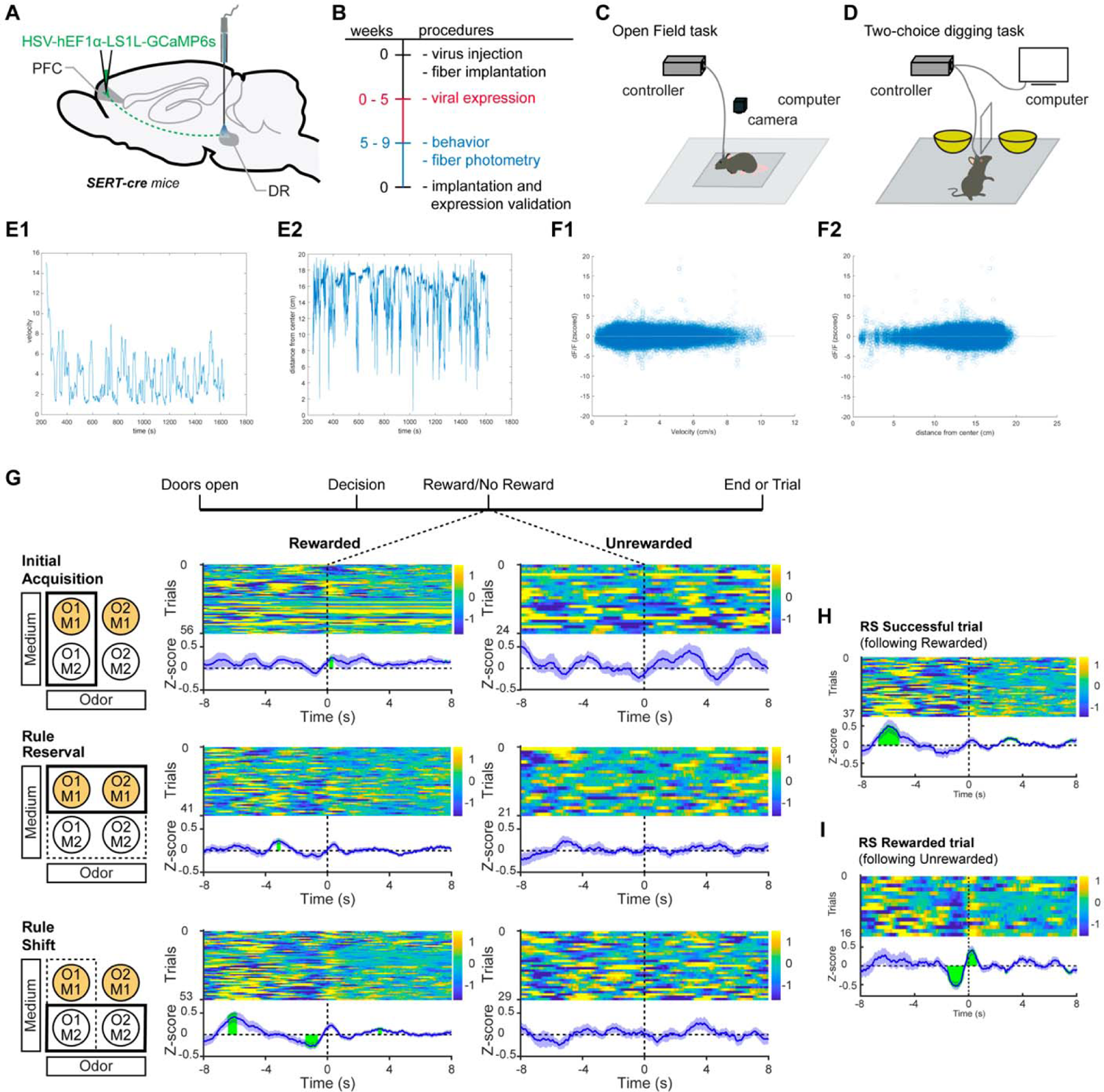
Serotonergic neurons projecting to the mPFC exhibit activity signatures during the extradimensional rule shift task. We injected HSV-hEF1α-LS1L-GCaMP6s into the mPFC of SERT-cre mice and recorded the activity of mPFC-projecting serotonergic neurons from the DR using fiber photometry (**A** and **B**), while mice were performing the open-field test (**C**) and the two-choice digging task (**D**). Examples for velocity (**E1**) and distance to center (**E2**) over time in the open field, where we observed no correlation between the activity of DR-to-mPFC serotonergic neurons and velocity (**F1**) or distance to the center (**F2**). In the two-choice digging task, we recorded neuronal activity during IA, RR, and RS and aligned traces to successful or unsuccessful reward retrieval. O = odor, M = medium. (**G**). For RS rewarded trials, we stratified data according to the outcome of the preceding trial (**H** and **I**). n = 4-6/group.

The open-field test assesses locomotor activity and anxiety-like behavior. We continuously recorded Ca^2+^ transients (100 Hz sampling frequency) along with position (10 Hz sampling frequency). No correlation was detected between serotonin neuronal activity in the DR (z-scored dF/F) and locomotion as assessed by velocity (R^2^ = 8.76×10^-5^, *P* = 0.0244, **Figure 2E1, F1**) or anxiety-like behavior as assessed by distance in center of the open-field test (R^2^ = 6.48×10^-5^, *P* = 0.0529, **Figure 2E2, F2**). Examples of individual correlations between DR serotonin neuronal activity and velocity and distance from the center are included in **Supplementary Figure 2A1** and **2A2**.

In the two-choice digging task, mice are challenged to adapt their behavior based on changes in rules which signal reward, a task that requires attention (Bissonette et al., 2013). Rules can change either within dimensions (intradimensional rule reversal, RR) or across dimensions (extradimensional rule shift, RS). In this test, we aligned DR serotonin neuronal activity to reward retrieval for rewarded trials and to the average latency of reward retrieval in rewarded trials for unrewarded trials. We measured neuronal activities during initial acquisition (IA), RR, and RS trials (**Figure 2G**).

For rewarded trials, activity is more dynamic pre-reward retrieval and less dynamic after reward retrieval (IA: *P* = 0.0017; RR: *P* = 4.100 ×10^-5^; RS: *P* = 3.653 ×10^-7^, **Figure 2G**). At reward retrieval we find a sharp biphasic response with reduced activity followed by a peak. For unrewarded trials, activity is dynamic throughout and no clear peaks or dips are discernable. For rewarded trials, activity signatures became more defined with increased task difficulty from IA to RR to RS. For rewarded RS trials, a clear signature emerged, with a broad peak at −6 s and the sharp signature around 0 s. Next, we further analyzed responses for rewarded RS trials by stratifying data based on whether the preceding trial was rewarded or unrewarded (**Figure 2H-I**). Interestingly, we detected the broad peak at −6 s only in trials following a successful trial (**Figure 2H**). Conversely, in trials following an unrewarded trial we detected a stronger biphasic response at reward retrieval (**Figure 2I**).

We also aligned DR serotonin neuronal activity to dig/choice and we did not detect significant changes in activity in ASST tasks (**Supplementary Figure 2B-G**).

Together these data demonstrate task-specific and dynamic activity signatures of serotonergic neurons projecting to the mPFC, with a role in cognitive flexibility, in particular extradimensional RS, but not for locomotor activity or anxiety-like behavior.

### Optogenetic inhibition of serotonergic terminals in the PL impairs extradimensional rule shift performance, without impacting exploratory or anxiety-like behavior

To determine whether endogenous PL serotonin is required for exploratory activity, anxiety-like behavior, and cognitive flexibility, we used an *in vivo* optogenetic approach, implanting optic fibers into the PL of a mouse line expressing the inhibitory *archaerhodopsin* (Arch) exclusively in serotonergic neurons, ePet1-Cre^-/+^::Ai35^+/+^ (Teixeira et al., 2018) (**Figure 3A-D, Supplementary Figure 7**).

**Figure 3.**
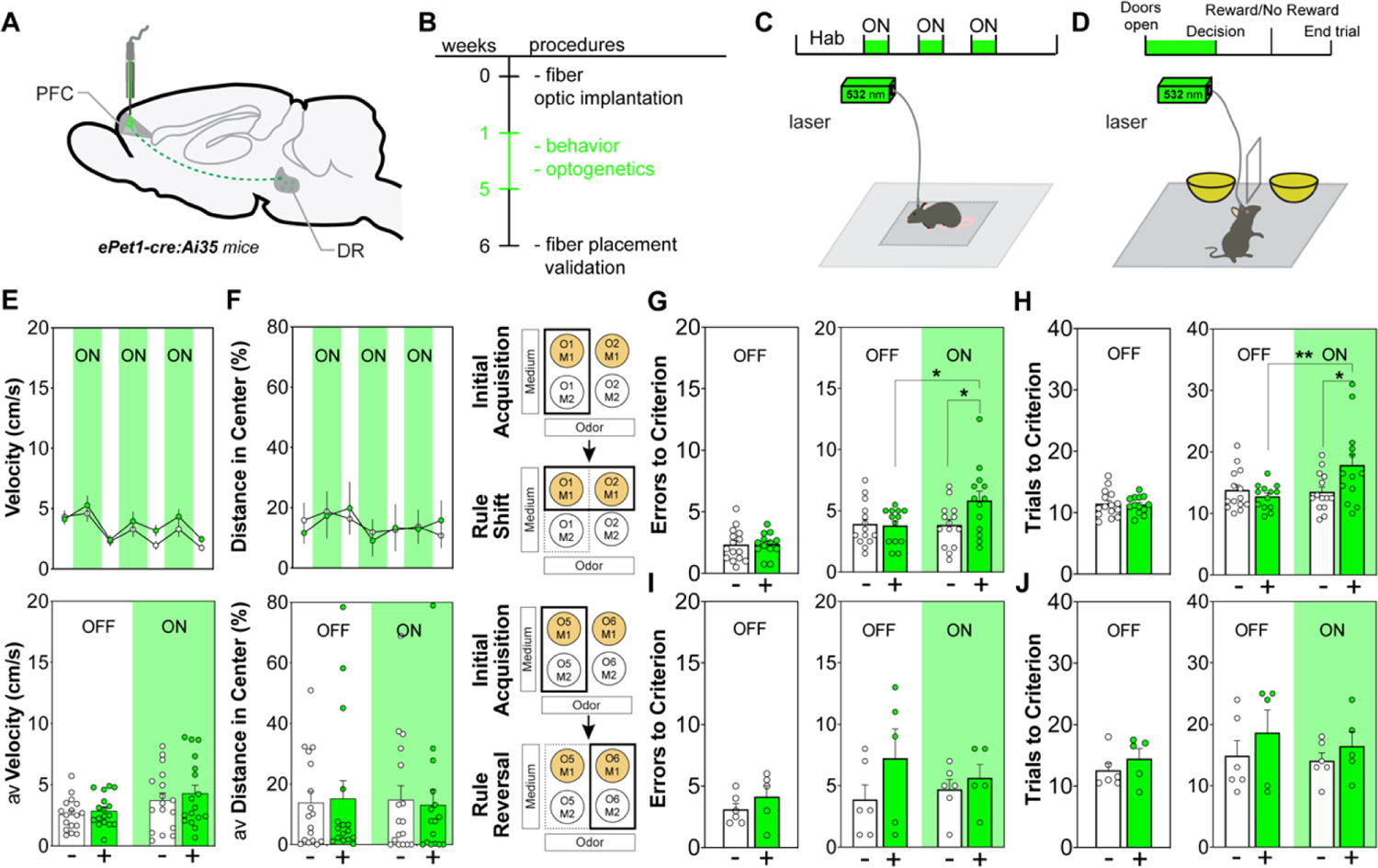
Optogenetic inhibition of serotonergic terminals in the PL impairs cognitive flexibility. We implanted ePet1-Cre^+^::Ai35^+/+^ (+) and ePet1-Cre^-^::Ai35^+/+^ (-) control mice in the PL for terminal opto-inhibition of serotonergic neurons projecting to the mPFC (**A** and **B**). Behavior was assessed in the open-field (**C**) and the two-choice digging test (**D**). No changes in velocity (**E**) and % distance in center (**F**) were detected in the open-field test. In the two-choice digging test, optogenetic inhibition of serotonergic terminals in the PL led to an increase in the number of errors and trials to criterion in the extradimensional rule shift task (**G** and **H**). Behavior in the intradimensional rule reversal task remained unaffected (**I** and **J**). Student’s paired and unpaired t-tests were used for posthoc comparisons. n = 13-18/group. *P < 0.05, ** P < 0.01.

In the open field test, we observed no impact of optogenetic inhibition in neither exploratory activity (OF_velocity_ = Genotype x Time interaction: across epochs, F_(6,186)_ = 1.084, *P* = 0.3734; Genotype x Stimulation interaction: all on and off epochs combined, F_(1, 32)_ = 0.0666, *P* = 0.7972; **Figure 3E** | OF_total distance_ = Genotype x Time interaction: across epochs, F_(6, 186)_ = 1.085, *P* = 0.3731; Genotype x Stimulation interaction: all on and off epochs combined, F_(1, 31)_ = 0.204, *P* = 0.6531; **Supplementary Figure 3A**) nor anxiety-like behavior (OF_distance center_ = Genotype x Time interaction: across epochs, F_(6, 186)_ = 0.4179, *P* = 0.8665; Genotype x Stimulation interaction: all on and off epochs combined, F_(1, 31)_ = 0.09184, *P* = 0.7629; **Figure 3F** | OF_time center_ = Genotype x Time interaction: across epochs, F_(6, 180)_ = 1.139, *P* = 0.3415; Genotype x Stimulation interaction: all on and off epochs combined, F_(1, 30)_ = 0.5313, *P* = 0.4889; **Supplementary Figure 3B**).

To assess the impact of reduced PL serotonin release on cognitive flexibility, we tested performance with and without opto-inhibition in the RS and RR tasks. We analyzed behavior for RS, because RS-performance is mPFC dependent (Dias et al., 1996a, Rogers et al., 2000) and serotonergic activity was strongest during this task (**Figure 4D**). We analyzed behavior for RR as a comparison, because RR-performance is thought to rely strongly on orbitofrontal cortex function and we found less mPFC serotonergic engagement. We furthermore focused our experimental manipulation on the pre-reward activity (broad peak at −6 s), because this was the most salient serotonergic activity signature specific to RS trials. Before each task, baseline learning was assessed during IA without opto-stimulation (Biro et al., 2019, Bissonette et al., 2008). At IA, prior to RS, and without light stimulation, ePet1-Cre^-/+^::Ai35^+/+^ and ePet1-Cre^-/-^::Ai35^+/+^ mice exhibited similar number of errors (Unpaired t-test: t_26_ = 0.02057, *P* = 0.9837; **Figure 3G**) and trials to criterion (Unpaired t-test: t_26_ = 0.3152, *P* = 0. 3776; **Figure 3H**). Upon light stimulation, ePet1-Cre^-/+^Ai35^+/+^ mice exhibited worse performance in the RS task than ePet1-Cre^-/-^::Ai35^+/+^. We detected a Genotype x Stimulation interaction in errors (two-way ANOVA: F_(1,25)_ = 4.061, *P* = 0.0493; **Figure 3G**) and trials (two-way ANOVA: F_(1,25)_ = 6.299, *P* = 0.0154; **Figure 3H**) to criterion. *Posthoc* comparisons showed no difference in significant differences between ePet1-Cre^-/-^::Ai35^+/+^ and ePet1-Cre^-/+^::Ai35^+/+^ in absence of light (errors to criterion: Sidak: t_50_ = 0.3371, *P* = 0.9311, **Figure 3G** | trials to criterion: Sidak: t_50_ = 0.7851, *P* = 0.682, **Figure 3H**). In presence of light, ePet1-Cre^-/+^::Ai35^+/+^ performed that task with significantly more errors (Sidak: t_50_ = 2.513, *P* = 0.0303, **Figure 3G**) and trials to criterion (Sidak: t_50_ = 2.764, *P* = 0.0159, **Figure 3H**) than ePet1-Cre^-/-^::Ai35^+/+^mice. Furthermore, *Posthoc* comparisons revealed an effect of stimulation in ePet1-Cre^-/+^::Ai35^+/+^ mice in errors (Sidak: t_50_ = 2.532, *P* = 0.0288, **Figure 3G**) and trials to criterion (Sidak: t_50_ = 3,173, *P* = 0.0052, **Figure 3H**).

**Figure 4.**
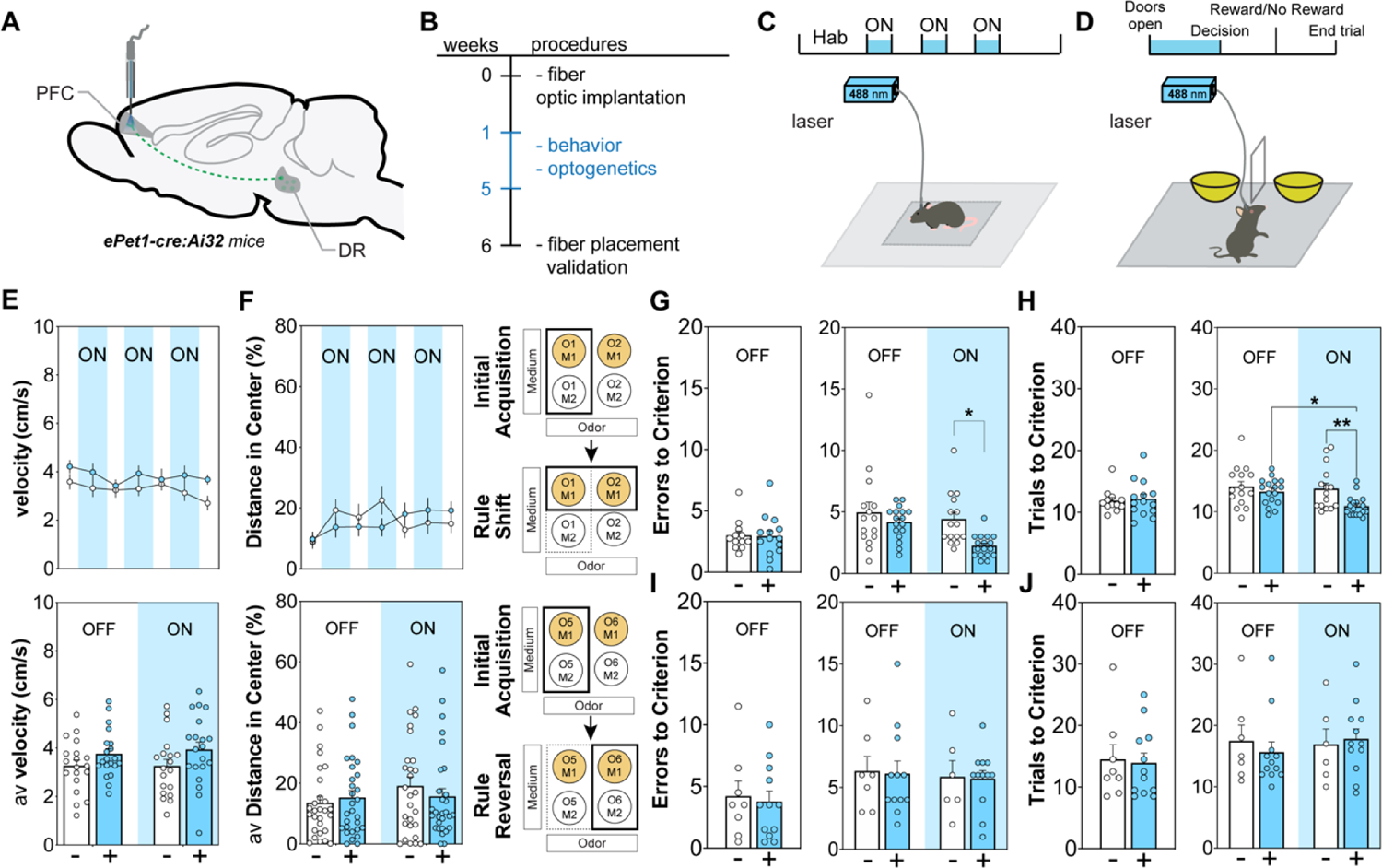
Optogenetic activation of serotonergic terminals in the PL increases cognitive flexibility. We implanted ePet1-Cre^+^::Ai32^+/+^ (+) and ePet1-Cre^-^::Ai32^+/+^ (-) control mice in the PL for terminal opto-stimulation of serotonergic neurons projecting to the mPFC (**A** and **B**). Behavior was assessed in the open-field (**C**) and the two-choice digging test (**D**). No changes in velocity (**E**) and % distance in center (**F**) were detected in the open-field test. In the two-choice digging test, optogenetic inhibition of serotonergic terminals in the PL led to a decrease in the number of errors and trials to criterion in the extradimensional rule shift task (**G** and **H**). Behavior in the intradimensional rule reversal task remained unaffected (**I** and **J**). Student’s paired and unpaired t-tests were used for posthoc comparisons. n = 12-21/group. *P < 0.05, ** P < 0.01.

Mice were then subjected to the RR task (**Figure 3I-J**). Prior to RR and in the absence of light, mice exhibited similar performance in IA (errors to criterion: Unpaired t-test: t_9_ = 1.020, *P* = 0.3346; **Figure 3I** | trials to criterion: Unpaired t-test: t_9_ = 0.9499, *P* = 0.3670; **Figure 3J**). For RR, we did not observe significant Genotype x Stimulation interactions in errors (F_(1,9)_= 0.9632, *P* = 0.3394; **Figure 3I**) or trials to criterion (F_(1,9)_= 0.1223, *P* = 0.7306; **Figure 3J**) between ePet1-Cre^-/+^::Ai35^+/+^ and ePet1-Cre^-/-^::Ai35^+/+^ mice. Both experimental groups exhibited similar latency to dig in both tasks (RS, Genotype x Stimulation interaction: two-way ANOVA: F_(1,25)_ = 0.00296, *P* = 0.9568; **Supplementary Figure 3C** | RR, Genotype x Stimulation interaction: two-way ANOVA: F_(1,9)_ = 0.1572, *P* = 0.6964; **Supplementary Figure 3D)**. Taken together, these data demonstrate that inhibition of serotonin release in the PL does not affect exploratory activity, anxiety-like behavior, or RR performance, but specifically impairs performance in the RS task of the two-choice digging task.

### Optogenetic stimulation of serotonergic terminals in the PL improves extradimensional rule shift performance, without impacting exploratory or anxiety-like behavior

Next, we investigate whether enhancing activity of serotonergic neurons projecting to the mPFC would alter exploratory activity, anxiety-like behavior, and cognitive flexibility. For that purpose, we used an *in vivo* optogenetic stimulation approach, implanting optic fibers into the PL of a mouse line expressing the excitatory channelrhodopsin-2 (ChR2) exclusively in serotonergic neurons, ePet1-Cre^-/+^::Ai32^+/+^ (Teixeira et al., 2018) (**Figure 4A, Supplementary Figure 7**).

In the open-field test, optogenetic stimulation of mPFC serotonergic terminals does not impact exploratory activity (OF_velocity_ = Genotype x Time interaction: across epochs, F_(6, 222)_ = 0.8700, *P* = 0.5177; Genotype x Time interaction: all on and off epochs combined, F_(1, 37)_ = 0.1146, *P* = 0.7359; **Figure 4E** | OF_total distance_ = Genotype x Time interaction: across epochs, F_(6, 330)_ = 0.5752, *P* = 0.75; Genotype x Time interaction: all on and off epochs combined, F_(1, 55)_ = 0.001575, *P* = 0.9684; **Supplementary Figure 4A**) and anxiety-like behavior (OF_distance center_ = Genotype x Time interaction: across epochs, F_(6, 330)_ = 2.494, *P* = 0.0225; Genotype x Stimulation interaction: all on and off epochs combined, F_(1, 55)_ = 0.985, *P* = 0.3232; **Figure 4F** | OF_time center_ = Genotype x Time interaction: across epochs, F_(6, 330)_ = 1.073, *P* = 0.3784; Genotype x Stimulation interaction: all on and off epochs combined, F_(1, 55)_ = 0.0817, *P* = 0.7755; **Supplementary Figure 4B**) Likewise, in the elevated-plus maze, optogenetic stimulation of mPFC serotonergic terminals does not impact optogenetic stimulation of mPFC serotonergic terminals does not impact anxiety-like behavior (**Supplementary Figure 5**).

Next, we assessed the effect of opto-stimulating the DR-to-PL serotonergic pathway on performance in the attention set-shifting tasks. As expected, during IA prior to RS, there was no effect of genotype on the number of errors (Unpaired t-test: t_24_ = 0.1129, *P* = 0.9110; **Figure 4G**) or the number of trials to criterion (Unpaired t-test: t_24_ = 0.3135, *P* = 0.7566; **Figure 4H**). In the RS task, selective photoactivation of PL serotonergic terminals reduced the number of errors (Genotype x Stimulation interaction: F_(1,30)_ = 1.751, *P* = 0.1908; main effect of genotype: two-way ANOVA: F_(1,30)_ = 8.023, *P* = 0.0063; **Figure 4G**) and trials to criterion (Genotype x Stimulation interaction: F_(1,30)_ = 2.185, *P* = 0.1446; main effect of genotype: two-way ANOVA: F_(1,30)_ = 7.296, *P* = 0.009; **Figure 4H**). *Posthoc* comparisons revealed no differences in performance in absence of light (errors to criterion: Sidak: t_60_ = 1.067, *P* = 0.4960, **Figure 4G** | trials to criterion: Sidak: t_60_ = 0.8487, *P* = 0.6393, **Figure 4H**). *Posthoc* comparisons also revealed that mPFC serotonergic photostimulation led to fewer errors (Sidak: t_60_ = 2.938, *P* = 0.0093, **Figure 4G**) and trials to criterion (Sidak: t_60_ = 2.955, *P* = 0.0089, **Figure 4H**) in ePet1-Cre^-/+^::Ai32^+/+^ mice in comparison to ePet1-Cre^-^::Ai32^+/+^ mice also exposed to light.

These observations demonstrate improved cognitive flexibility in the RS task as a result of optogenetic mPFC serotonergic terminal stimulation. Next, we assessed performance in the IA (baseline) and RR (with experimental manipulation). In IA, we observed similar performance of ePet1-Cre^-/+^::Ai32^+/+^ and ePet1-Cre^-/-^::Ai32^+/+^ mice in the absence of light, denoted by no differences in the number of errors (Unpaired t-test: t_18_ = 0.3582, *P* = 0.7243; **Figure 4I)** and trials to criterion (Unpaired t-test: t_18_ = 0.2981, *P* = 0.769; **Figure 4J**). In the RR task (**Figure 4I-J**), we did not observe significant Genotype x Stimulation interactions in errors (F_(1,16)_= 0.0003, *P* = 0.987; **Figure 4I**) or trials to criterion (F_(1,16)_= 0.5977, *P* = 0.445; **Figure 4J**) between ePet1-Cre^-/+^::Ai32^+/+^ and ePet1-Cre^-/-^::Ai32^+/+^ mice. Both experimental groups exhibited similar latency to dig in both tasks (RS, Genotype x Stimulation interaction: two-way ANOVA: F_(1,14)_ = 0.3457, *P* = 0.5613; **Supplementary Figure 6A** | RR, Genotype x Stimulation interaction: two-way ANOVA: F_(1,36)_ = 0.57, *P* = 0.4552; **Supplementary Figure 6B)**. Taken together, these results demonstrate that increased PL serotonergic signaling improves performance in the two choice digging task, specifically in RS trials, but not RR trials.

## Discussion

We find that serotonin in the PL modulates cognitive flexibility, specifically extradimensional rule shifting performance. Using fiber photometry we demonstrate that DR serotonergic neurons that project to the PL sub-region of the mPFC are most active prior to reward retrieval in the extradimensional RS task of the two-choice digging test. Using optogenetics we show that a decrease of terminal serotonergic activity in the mPFC during the RS task impairs performance, while an increase leads to improved performance. During intradimensional RR, endogenous serotonergic activity is weaker and functional manipulations of serotonergic terminal activity in the mPFC have no impact on behavioral performance. Our data thus demonstrate that the activity of DR-to-mPFC serotonergic neurons is necessary for normal, and sufficient to improve, cognitive flexibility for extradimensional RS but not intradimensional RR performance.

In electrophysiological experiments, we demonstrated a primary and direct pyramidal inhibition of serotonin in the LV of the PL subregion through a 5-HT_1A_ receptor-mediated mechanism. *Ex vivo* optogenetic stimulation of mPFC serotonergic terminals suppresses spike firing of local LV pyramidal neurons, an inhibitory action blocked by 5-HT_1A_ receptor antagonist WAY100635. The observed outward currents in voltage-clamp and cessation of spiking in current clamp are consistent with somato/dendritic 5-HT_1A_ receptor activation of G protein-gated inwardly rectifying potassium (GIRK) channels (Anderson et al., 2006, Rojas et al., 2016) as well as 5-HT_1A_ receptor-mediated suppression of voltage-gated sodium channels at the axon initial segment (Yin et al., 2017). The observed inhibition was sensitive to 5-HT_1A_ receptor blockade, consistent with the previously-reported effect of serotonin opto-stimulation on LVI pyramidal neurons (Sparks et al., 2017). Here, we demonstrate that this inhibition of the major output neurons is unperturbed by synaptic blockers. In transgenic ePet1-Cre^+/-^::Ai32^+/+^ mice, terminal serotonin exerts an acute inhibitory 5-HT_1A_ receptor-mediated control over deep layer mPFC pyramidal neurons.

Using fiber photometry, we identified an activity signature of mPFC-projecting DR serotonergic neurons specifically during rewarded extradimensional RS trials: an elevation of Ca^2+^ transients prior to reward retrieval (around −6 s) and a sharp biphasic response at reward retrieval. This general finding is in line with results from several studies in rodents and monkeys that demonstrate increased firing activity of DR neurons, in some cases identified as serotonergic, in response to reward and reward-predicting cues (Bromberg-Martin et al., 2010, Cohen et al., 2015, Inaba et al., 2013, Nakamura et al., 2008, Ranade et al., 2009). Because we did not deliver discrete cues with high temporal precision the peak we observe at around −6 s could relate to more timely diffuse cue-perception. In contrast to prior studies, we selectively analyzed serotonergic DR-to-mPFC pathway-specific population responses. Interestingly, the activity signature scaled and became more pronounced from IA to RR to RS. Thus, the activity signature scales with mPFC dependence of the task and is in line with the functional dissociation between the mPFC and OFC in RS and RR, respectively (Birrell et al., 2000, Bissonette et al., 2015, Dias et al., 1996a, Dias et al., 1996b, Fellows et al., 2003, Kringelbach et al., 2003, O’Doherty et al., 2003, Ragozzino, 2007, Rogers et al., 2000, Rygula et al., 2010). The population response at reward retrieval when following a failed (non-rewarded) trial is larger than when following a successful (reward) trial. This finding is congruent with serotonergic neurons showing a larger response to unexpected versus expected rewards (Bromberg-Martin et al., 2010, Cohen et al., 2015, Grossman et al., 2022, Inaba et al., 2013, Nakamura et al., 2008, Ranade et al., 2009). The broad peak at −6 s was significant for rewarded trials following a successful (rewarded) trial but not following a failed (non-rewarded) trial. Thus a combination of cues that were positively reinforced in the prior trial might become more predictive in the subsequent trial and trigger a larger serotonergic response. This interpretation is in line with the reported activity of DR serotonergic neurons to reward-predicting cues (Bromberg-Martin et al., 2010, Cohen et al., 2015, Inaba et al., 2013, Nakamura et al., 2008, Ranade et al., 2009). Because the ASST assesses cognitive flexibility in the context of multidimensional reward-predicting cue associations, we tested the role of the potentially cue-related activity around −6 s in behavioral performance.

We found that optogenetic inhibition of serotonergic DR-to-mPFC input from trial start to dig, which covers the period of increased serotonergic activity during rewarded RS trials, impairs performance in the extradimensional RS task. Conversely, optogenetic activation of serotonergic DR-to-mPFC input during the same period improves performance in the extradimensional RS task. This finding is broadly in line with chemogenetic inhibition of DRN 5-HT neurons causing perseverative responding (Matias et el., 2017). We furthermore found that neither optogenetic inhibition nor excitation impacted RR performance. This finding is in contrast to the inhibition of DR serotonergic neurons slowing reversal learning (Matias et el., 2017). However, both findings can be easily reconciled by the fact that Matias et al manipulated DR serotonergic neurons, while we selectively manipulated DR serotonergic neurons that project to the PL. Hence we predict that DR serotonergic neuron inhibition captures the population of neurons that project to the mPFC and therefore also impairs RS performance. The selective specificity for RS performance is congruent with our findings of increased endogenous serotonergic activity during the RS task when compared to the RR task. Our findings are also again in line with the functional dissociation between the mPFC and OFC in RS and RR, respectively (Birrell et al., 2000, Bissonette et al., 2015, Chudasama et al., 2003, Dias et al., 1996a, Dias et al., 1996b, Fellows et al., 2003, Kringelbach et al., 2003, O’Doherty et al., 2003, Ragozzino, 2007, Rogers et al., 2000, Rygula et al., 2010). Most strikingly, OFC serotonin depletion in marmosets impairs reversal learning (Clarke et al., 2004), without affecting set-shifting (Clarke et al., 2005). Taken together, these findings indicate that patterns and rules we identified for serotonergic input into the mPFC during RS are also present for serotonergic input into the OFC during RR. Future studies will test this hypothesis.

In low- and moderate-threat environments such as the open field, dorsal raphe serotonergic neurons display reduced activity with movement, and opto-stimulation of DR serotonergic neurons suppresses movement (Seo et al., 2019). We find no correlation between the activity of serotonergic neurons projecting from the DR to the mPFC, and no effect of opto-stimulation of this projection on locomotion in the open field. This finding indicates that DR serotonergic neurons that code for and modulate movement do not project to the mPFC.

Opto-stimulation of DR serotonergic neurons also increases ability to wait for delayed rewards or patience (Fonseca et al., 2015, Lottem et al., 2018, Miyazaki et al., 2018, Miyazaki et al., 2014). In contrast, we do not find effects of opto-stimulation or inhibition on the latency to dig in the ASST, indicating no effect on impulsivity/patience. However, our experiments did not assess the effect of delayed rewards and we can therefore not draw direct comparisons between studies. It will be interesting to see if RS performance paired with delayed reward delivery will further increase endogenous serotonergic activity as well as the effect size of inhibition or activation of serotonergic input into the PL on behavioral performance.

The mPFC is also associated with emotional regulation (Anderson et al., 2006, Barrash et al., 2000, Bechara et al., 1994, Damasio et al., 1990, Kroes et al., 2019, Szczepanski et al., 2014). However, we find no evidence for a role of serotonin input into the mPFC in emotional regulation. Specifically, we find no GCaMP-based activity signatures that relate to behavior in the open field test, nor do we find an effect on behavior in the open field test caused by optogenetic inhibition or excitation. Our findings are in contrast to the anxiolytic effect of exogenous serotonin and 5-HT_1A_ receptor agonist (Yamashita et al., 2018), as well as 5-HT_1B_ receptor agonist (Kjaerby et al., 2016) infusions into the mPFC. One likely explanation for this discrepancy lies in the spatio-temporal differences of receptor engagement between the two approaches. While optogenetically triggered terminally released serotonin preferentially binds to receptors in close proximity to the release site, intra-PL injected serotonin and serotonin receptor agonists likely flood the entire extracellular space and activate receptors indiscriminately of location relative to endogenous release sites. Serotonergic projections to the mPFC mainly arise from the rostral and central subregions of the DR (Chandler et al., 2013, Fernandez et al., 2016, Muzerelle et al., 2016, Prouty et al., 2017). Thus, findings that optogenetic and pharmacogenetic manipulations of DR serotonergic neurons do not change anxiety-like behavior (Correia et al., 2017, Ohmura et al., 2014) (Teissier et al., 2015) while MR manipulations do (Ohmura et al., 2014, Teissier et al., 2015) are congruent with our data.

Collectively, our data demonstrate a powerful role for serotonin input into the mPFC in cognitive flexibility. Many psychiatric disorders present with deficits in cognitive flexibility, for example ruminations in depression, compulsion in substance use disorders and obsessive compulsive disorder, and the inability to form new associations in post-traumatic stress disorder. We speculate that serotonergic prescription drugs (such as serotonin reuptake inhibitors) and recreational drugs (such as psychedelics) may improve cognitive flexibility by interacting with serotonin signaling in the mPFC (Davis et al., 2020, Dos Santos et al., 2018, Manzano-Nieves et al., 2021, Murphy-Beiner et al., 2020, Torrado Pacheco et al., 2023, Winkelman, 2014). Our data provide a mechanistic foundation to investigate such relationships which will ultimately improve diagnosis and treatment approaches in psychiatry.

## Materials and Methods

### Animals

Adult mice aged 12-32 weeks were used for all behavioral studies, Mice were bred at Columbia Psychiatry, New York State Psychiatric Institute, group-housed and provided with food and water *ad libitum*, and were maintained on a 12-hour light/dark cycle (lights on at 7:00 AM), relative humidity of 55%, 22°C. Procedures were approved by the Animal Care and Use Committee of the New York State Psychiatric Institute and in accordance with the National Institute of Health (NIH) Guide for the Care and Use of Laboratory Animals.

Male Sert-cre mice (129S2/129SvEv/Tac or B6129SF1) mice were used for fiber photometry experiments and male ePet1-Cre::Ai32 F1 hybrid background (C57BL/6J x 129S2/129SvEv/Tac or B6129SF1) and ePet1-Cre::Ai35 (129SvEv) for optogenetic experiments. Transgenic mice expressing the light-gated cation *channel-rhodopsin-2* (ChR2) were generated as previously described (Abela et al., 2020, Browne et al., 2019, Teixeira et al., 2018). Next, mice from the serotonergic cre-driver line ePet1 ((Scott et al., 2005), Jackson 012712, Bar Harbor, Maine, USA) were crossed with the Cre-dependent ChR2 reporter line Ai32 (Ai32; Jackson 012569) for the direct serotonin-specific expression of ChR2. The line ePet1 was originally generated on the F2 background (C57BL/6 x SJL) and has been backcrossed in the lab to 129SvEv/Tac for >10 generations. The line Ai32 was generated on the F1 background (129S6/SvEvTac x C57BL/6NCrl) and has been backcrossed in our lab to 129SvEv/Tac for >10 generations. The Cre-dependent *archaerhodopsin* (Arch) reporter line (Ai35; Jackson 012735) was used for the specific expression of Arch in serotonergic neurons. Experimental mice were homozygous for Ai32 or Ai35 and positive for ePet1-cre ((ePet1-Cre^+/−^::Ai32^+/+^ (ChR2) or (ePet1-Cre^+/−^::Ai35^+/+^ (Arch)), while control mice were homozygous for Ai32 or Ai35 but negative for ePet-Cre ((ePet1-Cre^−/−^::Ai32^+/+^ (ChR2^-^) and ePet1-Cre^−/−^::Ai35^+/+^ (Arch^-^).

### Electrophysiological Recording and Analysis

Electrophysiology experiments were performed in acute brain slices obtained from male and female adult ePet1Cre^+^::Ai32^+/+^ mice and control ePet1-Cre^−^::Ai32^+/+^ mice. After deep anesthesia with chloral hydrate (400 mg/kg), mice were decapitated and their brains were quickly extracted and chilled in 4°C sucrose artificial cerebrospinal fluid (aCSF) (254 mM sucrose, 10 mM D-glucose, 24 mM NaHCO_3_, 2 mM CaCl_2_, 2 mM MgSO_4_, 3 mM KCl, 1.25 mM NaH_2_PO_4_; pH 7.4). A Dosaka linear slicer was used to obtain 400 μM coronal brain slices of PFC (range 2.34–0.74 from Bregma; Paxinos and Franklin, 2001), which recovered for ≤ 2 h in regular aCSF (128 mM NaCl, 10 mM D-glucose, 26 mM NaHCO_3_, 2 mM CaCl_2_, 2 mM MgSO_4_, 3 mM KCl, 1.25 mM NaH_2_PO_4_; pH 7.4). To maintain the synthesis of serotonin (Liu et al., 2005), brain slices were recovered and recorded in the presence of 30 µM L-tryptophan. Electrophysiological recordings were performed as described previously (Sargin et al., 2020). In brief, brain slices were placed in a perfusion chamber on the stage of a BX50W1 microscope (Olympus) and perfused with oxygenated aCSF (95%O_2_, 5%CO_2_) at a rate of 3–4 ml/minute at 30°C for cortical slices. In the PL and cingulate regions of the mPFC, LV pyramidal neurons were identified based on morphological characteristics (pyramidal shape, large cell body, orientation of apical dendrite) and patched using the following intracellular solution: 120 potassium gluconate, 5 mM KCl, 2 mM MgCl_2_, 4 mM K_2_-ATP, 0.4 mM Na_2_-GTP, 10 mM Na_2_-phosphocreatine and 10 mM HEPES buffer and with pH adjusted to 7.3 using KOH. Intrinsic properties of the recorded neurons were as follows: resting membrane potential: −78 ± 1 mV; input resistance: 115 ± 13 M-ohm; capacitance: 155 ± 10 pF; spike amplitude: 70 ± 3 mV (n = 70 neurons; 15 mice). Optogenetic stimulation was delivered by a blue light using a microscope-mounted collimated LED (470 nm, Thorlabs, Newton, New Jersey, United States) as follows: 10 ms light pulses in a 20 Hz train for 5 s, with a subset of neurons also receiving longer stimuli. Optogenetic stimulus episodes were separated by ≥ 5-min. Neurons were either held in voltage-clamp at −75 mV or in current-clamp with positive current injected to elicit ∼3 Hz action potential firing at baseline. Recordings were obtained using a Multiclamp 700 amplifier (Molecular Devices, San Jose, California, United States). All data were acquired at 20 kHz and low pass filtered at 3 kHz using pClamp software (Molecular Devices) and corrected for the liquid junction potential (14 mV). Pharmacological manipulations include CNQX (20 µM; Alomone Labs) and APV (50 µM; Alomone Labs) that were used to block glutamate receptors, bicuculline (3 µM; Tocris) that was used to block GABA receptors, and WAY100635 (30 nM; Tocris) that was used to block 5-HT_1A_ receptors.

### Viruses

HSV-hEF1α-LS1L-GFP and HSV-hEF1α-LS1L-GCaMP6s were purchased from the Massachusetts General Hospital Neuroscience Center vector core. HSV titer was 1.00 x 10^9^ genomic copies *per* ml.

### Stereotaxic Surgery

Mice undergo surgery for insertion of fiber optic fibers or viral injections, under isoflurane anesthesia (1.5-2-0% isoflurane gas/oxygen mixture (SomnoSuite, Kent Scientific, Torrington, Connecticut, United States) and using stereotaxic apparatus (Stoelting, Wood Dale, Illinois, United States). Throughout the entire surgery, body temperature was maintained with an adjustable warming pad. After surgical procedures, mice were treated with Carprofen (MediGel CPF, ClearH_2_O, Portland, Maine, United States) or Rimadyl (Bio-Serv, Flemington, New Jersey, United States) for pain management for three days.

For retrograde labeling, 150 nL of HSV-hEF1α-LS1L-GFP was bilaterally injected into mPFC (anterior-posterior, AP: +1.8; medial-lateral, ML: ± 0.3; dorsal-ventral, DV: −1.85 and − 2.0). Brains were collected 4 weeks later.

For *in vivo* optogenetics, fiber optic cannulae (1.25 mm white ceramic ferrule, 200 µm core 0.37 NA, Newdoon Technologies Co., Ltd, Hangzhou, China) consisting of 1.25 mm white ceramic ferrules with 200 µm optical fibers (Newdoon Technologies) were bilaterally implanted, targeting LII/III of the PL (one hemisphere: AP: +1.8; ML: + 0.3; DV: −1.8; and another hemisphere: AP: +1.8; ML: +0.8; DV −1.87 at 16° angle). Four weeks later when the viral expression stabilized, imaging was performed during the behavioral assessment.

For fiber photometry, 150 nL of HSV-hEF1α-LS1L-GCaMP6s was bilaterally injected into mPFC (AP: +1.8; ML: ± 0.3; −1.85 and −2.0) and a fiber optic cannula (2.5 mm flat tip metal ferrule, 0.48 NA, Doric, Quebec, Canada) implanted into DR (AP: +5.2, ML: +0.8; DV: −3.3 at 16°). Behavioral experiments started four weeks post-surgery.

Viruses were injected using Nanoject III (Drummond Scientific, Broomall, Pennsylvania, United States). Fiber optic cannulae were fixed to the skull using dental cement (Stoelting) mixed with carbon (Sigma-Aldrich, St. Louis, Missouri, United States).

### Behavioral Testing

Mice were subjected to the two-choice digging task and open-field, after recovering from surgery. All animals were exposed to the same behavioral paradigms and timeline. One week before behavior testing, mice were handled and tethered to optical patch cables daily for at least 5 minutes *per* mouse for 3 consecutive days. The behavioral assessment took place during the light cycle between 10 AM and 7 PM. To eliminate odor irrelevant cues, each apparatus was thoroughly cleaned with ethanol 10% after each animal.

#### Open-field test

The open-field test (OF) was performed to assess anxiety-like behavior, exploratory and locomotor activity. The OF apparatus consists of an open and transparent square arena (length and width, 40.7 cm) with transparent Plexiglas walls (30.5 cm high) equipped with infrared detectors to track animal movement (model ENV-520, Med Associates, Vermont, United States). Mice were placed in the center of the OF arena and allowed to freely explore it. Mouse’s movements were detected through infrared beams and the number of beams crossed was automatically recorded through the Kinder scientific software (Kinder scientific, Chula Vista, California, US). Velocity and the amount of time and distance traveled in the center (defined as the central 15 x 15 cm region) and periphery zone was recorded and analyzed.

#### Serial Rule Shifting Test (Two-Choice Digging Task)

The serial rule shifting test was performed as previously described by (Bissonette et al., 2008, Canetta et al., 2016, Cho et al., 2015, Goodwill et al., 2018). During the test, mice were food restricted (1.8g/mouse for 5 days and 2g/mouse thereafter), to maintain no less than 85% of free-feeding body weight. After at least 5 days of food restriction, mice were initially habituated and trained to dig in small ceramic bowls (3” Clay Saucers by Ashland®, Michaels Stores, Inc., Texas, United States) filled with bedding material (VitaKraft® World Bedding Grey and The Andersons® Bed-o’Cobs ¼, W. F. Fisher and Son, Inc, Branchburg, New Jersey, United States) to retrieve a food reward (bait is 1⁄4 of a honey nut loop (Honey Nut Cheerios©, General Mills, Golden Valley, Minnesota, United States) in their home cage overnight. On the next day, mice were habituated to the apparatus and ceramic bowls for 3 trials. In each trial, more bedding material was added to encourage digging behavior and reward more pronounced digging behavior in the bowls. The apparatus was an acrylic box (length, 31 cm; width, 21 or 20; height, 21 or 20 cm) comprised of a large waiting compartment (length, 20 cm; width, 29 cm) and two equally sized choice compartments (left (L) and right (R) – length, 17 cm; width, 10 cm) that could be accessed when opening the inserted sliding doors. Mice were initially placed in the larger compartment and each trial was initiated when the doors are lifted. After habituation, mice were trained to dig to obtain the reward on either side of the apparatus. Mice were trained in one session of 5 trials per day. Each mouse digs in each of the four exemplar pairs at least once. (i.e., on the left (L) and right (R): Bedding 1 (L) & Bedding 2 (R); 2 (L) & 1 (R); 1 (L) & 1 (R); and 2 (L) & 2 (R).

Following two successful training days (dig and collect the reward from both bowls in under 2 minutes), mice are ready to be tested. Mice learn to discriminate between four combinations of two odors and two bedding materials. All odors were ground dried spices (cinnamon and paprika, McCormick & Company, Baltimore, Maryland, United States) and unscented digging material. See **Supplementary Table 1** for reference on sequences of odor and texture associations. During IA and in the absence of light, mice learn that the reward is solely associated with a given odor (scent) and texture (bedding material) being subjected to a choice to dig between a correct (bowl containing the reward) and an incorrect (bowl with no reward) choice. Once mice achieve the criterion (8 correct trials out of 10), they were challenged with a rule change that leads to a different association. The change was either interdimensional (rule reversal (RR); e.g., odor A to odor B or digging material 1 to digging material 2) or extradimensional (rule shift (RS); e.g., odor A to digging media 1 or digging media 2 to odor B). During either IA and rule change tasks, the criterion was the same. Mice subjected to two-choice digging tasks performed up to two days of serial RR and/or four days of serial RS. Experimental and control mice were randomly divided into two groups, one group that received optogenetic stimulation or inhibition on day 1 (& 4), and another that received it on day 2 (& 3), in a counterbalanced manner (ON-OFF(-OFF-ON)) or OFF-ON(-ON-OFF) sequence). This experimental design accounts for an order and/or sequence effect. Counterbalancing was performed for both RS (four days) and RR (two days). In all experimental groups, IA was performed in absence of optogenetic manipulation. The number of errors, trials, and latency was recorded. Mice had up to 50 trials to achieve the criterion. Each trial was limited to 8 minutes. The average intertrial interval (ITI) for FP experiments across stages was 160.37 seconds (SEM = 13.29 s): IA: mean= 179.19 s (SEM = 16.63 s); RR: mean = 162.4 s (SEM = 12.99 s); and RS: mean = 139.52 (SEM = 10.24 s). Sessions ended when they either after three times out, or the criterion was met. See Table S1 for reference on sequences of odor and texture associations.

#### Fiber Photometry

Ca^2+^ signal fluctuations were recorded in Sert-cre mice implanted with fiber optic cannulae into the DR, using a RZ5P processor, and captured using the Synapse software (Tucker-Davis Technologies (TDT), Alachua, Florida, United States). Signals will be acquired at 1013 Hz with LED transmitting 30 μW of light on average. The LEDs used were 405 nm blue, as a Ca^2+^-independent isosbestic control signal for motion and photobleaching, and 465 nm purple, as a Ca^2+^-dependent signal (Doric). To quantify changes in fluorescence, deltaF/F, a least-squares linear fit was applied to the 405 nm signal and aligned to the 465 nm signal, producing a fitted 405 nm signal to normalize the 465 nm signal as follows: deltaF/F = (465 nm signal – fitted 405 nm signal)/fitted 405 nm signal. The imaging and behavioral files were analyzed using customized scripts on MATLAB (MathWorks, Natick, Massachusetts, United States). Briefly, we applied data processing steps previously described in the literature: a global detrending method (Liang et al., 2018) a high-pass filter (Cho et al., 2017), and zeroing to the 8^th^ percentile to our deltaF/F signal (Jones et al., 2020). The signal was then transformed into a robust z-score for comparison across animals. During OF trials fiber photometry recordings lasted for 35 minutes. In the two-choice digging test, fiber photometry recordings occurred throughout the whole trials, successful and unsuccessful, of IA, RS, and RS tasks. We then aligned the behavioral trace to the imaging trace and looked at the z-scored values within time windows that spanned behavioral transitions, both at the moment of the dig/choice and the reward retrieval.

#### In vivo optogenetics

Mice were tethered to optical patch cables coupled to an optical commutator (Doric) and connected to a 473 nm blue light and 532 nm green light diode-pumped solid-state laser (Laserglow Technologies, Ontario, Canada). Laser output was controlled by a waveform generator (Master 8 or Master 9, A.M.P.I., Jerusalem, Israel). An Arduino UNO controlled, four channels, simultaneous TTL pulse generator, and controller with BNC connectors can be used in lieu of the Master 8 or 9 for pulse control. ePet1Cre+::Ai32+/+ and ePet1-Cre−::Ai32+/+ mice were stimulated using blue light pulses (20 Hz, 10 ms, 8-10 mW) while ePet1Cre+::Ai35+/+ and ePet1-Cre−::Ai35+/+ mice were stimulated with green light (constant, 8-10 mW) continuously for up to 50 seconds. In the open-field test, optogenetic stimulation was presented for three cycles of 3 minutes light ON periods interspersed with two 3 minutes light OFF periods. Optogenetic inhibition was presented in three cycles of 50 seconds of light ON interspersed with two 3 minutes light OFF periods. For both stimulation and inhibition experiments, a period of ∼10 minutes in absence of light was included, prior, and after light ON/OFF cycles. Experiments that included photostimulation and photoinhibition lasted for a total of 35 and 28.5 minutes, respectively. In optogenetic experiments, while performing the two-choice digging test, stimulation or inhibition occurred during each trial from the start until the mouse digs in one of the two bowls (correct or incorrect) and lasted up to 8 minutes (stimulation) or 50 seconds (inhibition).

#### Immunofluorescence

For YFP, serotonin, and 5-HTT immunofluorescence procedures, mice were perfused transcardially with 4°C 0.1 M phosphate-buffered saline (PBS) and then 4% paraformaldehyde (PFA). Brains were removed, fixed overnight in PFA, and then transferred to a 30% sucrose solution and stored at 4°C. PFC and DR regions were sectioned using a vibratome and 50mm coronal free-floating sections were prepared for immunofluorescence. For immunodetection, sections were initially blocked for 2 hours at room temperature (RT) with PBS-T 0.05% containing 10% of a donkey or goat Serum (Sigma). Primary antibodies, anti-GFP (chicken, 1:5000, ab13970, Abcam, Cambridge, United Kingdom), Anti-serotonin (goat, 1:500, ab66047, Abcam), anti-5-HTT (guinea pig, 1:50, HTT-GP-Af1400, Nittobo Medical Co, Japan) and anti-NeuN (rabbit, 1:200, MABN140, Millipore, Burlington, Massachusetts, United States) were incubated overnight at RT with PBS-T 0.3% with 2% donkey or goat serum. Then, sections were incubated for 2 hours at RT with secondary antibodies (488 donkey anti-rabbit, 555 or 568 donkey anti-goat; 1:1000; Thermo Fischer Scientific, Waltham, Massachusetts, United States). Hoechst was used to stain cell nuclei. Sections were mounted in ProLong (ThermoFisher scientific, Waltham, Massachusetts, United States) mounting medium.

### Confocal Microscopy

Imaging was performed on a laser confocal microscope (TCS SP8, Leica Microsystems, Wetzlar, Germany) operated and visualized through the LAS X software v 3.5.5.19976 (Leica). Images were acquired with a 20x objective. Detection settings were set for each laser: “far red” (638 nm), “red” (552 nm), “green” (488 nm), and “blue” (405 nm). All image stacks were acquired with a stepped thickness of 2 μm and z-stack compilation was performed using either the LAS X or Fiji (ImageJ) software.

### Statistical analysis

Statistical analysis for behavior was performed using Prism 9.4.0 software (GraphPad, La Jolla, California, United States). Data were analyzed using a two-tailed Student’s t-test, to compare the mean values for two groups, for comparisons between on and off periods, two-way ANOVA, and for longitudinal analyses a repeated measures ANOVA, with Sidak posthoc testing as indicated. The criterion for significance for all analyses was set at *P < 0.05; **P < 0.01; ***P < 0.001. Fiber photometry data were analyzed with custom-written MATLAB scripts. The 405 nm control trace was fitted to and subtracted from the 470 nm trace to calculate a dF, which was then divided by the fitted control trace to calculate a dF/F. The dF/F was globally detrended, high-pass filtered, and zero corrected to the 8^th^ percentile using a moving window. The processed dF/F traces were converted to z-scores based on their median absolute deviation from the median and then smoothed. For the cognitive flexibility analysis, the traces were aligned to behaviorally coded events. The aligned traces were non-normally distributed based on the Kolmogorov-Smirnov test, so we used a Wilcoxon rank sum test to find significant time points with P < 0.005. For the open field analysis, each fully processed dF/F value was plotted against either its corresponding distance-from-center position or velocity for each time frame. A least-squares line was plotted, and the corresponding statistics were reported. The effect size was estimated from previous experiments. Results are expressed as mean ± standard error of the mean (SEM). Statistical details were included in Table 2.

## Acknowledgements

We gratefully acknowledge the contributions of Simone Valade and Yao-Fang Tan to the electrophysiological experiments. We also thank Marie Labouesse and Arturo Torres-Herraez for their assistance with the analysis of fiber photometry data. This work was supported by Columbia University (Innovation Award, NDA), NSF (GRFP DGE16-44869, AAM), NIMH (R01MH113569, 2R01MH080116, MSA), an NSERC Discovery Grant (EKL), and a CIHR Canada Graduate Scholarship Doctoral Award (SKP).

## References

Abela AR, Browne CJ, Sargin D, Prevot TD, Ji XD, Li Z, Lambe EK, Fletcher PJ (2020) Median raphe serotonin neurons promote anxiety-like behavior via inputs to the dorsal hippocampus. Neuropharmacology 168: 107985

Adhikari A, Topiwala MA, Gordon JA (2011) Single units in the medial prefrontal cortex with anxiety-related firing patterns are preferentially influenced by ventral hippocampal activity. Neuron 71: 898–910

Amargos-Bosch M, Bortolozzi A, Puig MV, Serrats J, Adell A, Celada P, Toth M, Mengod G, Artigas F (2004) Co-expression and in vivo interaction of serotonin1A and serotonin2A receptors in pyramidal neurons of prefrontal cortex. Cereb Cortex 14: 281–99

Anderson SW, Barrash J, Bechara A, Tranel D (2006) Impairments of emotion and real-world complex behavior following childhood- or adult-onset damage to ventromedial prefrontal cortex. J Int Neuropsychol Soc 12: 224–35

Barrash J, Tranel D, Anderson SW (2000) Acquired personality disturbances associated with bilateral damage to the ventromedial prefrontal region. Dev Neuropsychol 18: 355–81

Bechara A, Damasio AR, Damasio H, Anderson SW (1994) Insensitivity to future consequences following damage to human prefrontal cortex. Cognition 50: 7–15

Biro S, Lasztoczi B, Klausberger T (2019) A Visual Two-Choice Rule-Switch Task for Head-Fixed Mice. Front Behav Neurosci 13: 119

Birrell JM, Brown VJ (2000) Medial frontal cortex mediates perceptual attentional set shifting in the rat. J Neurosci 20: 4320–4

Bissonette GB, Martins GJ, Franz TM, Harper ES, Schoenbaum G, Powell EM (2008) Double dissociation of the effects of medial and orbital prefrontal cortical lesions on attentional and affective shifts in mice. J Neurosci 28: 11124–30

Bissonette GB, Powell EM, Roesch MR (2013) Neural structures underlying set-shifting: roles of medial prefrontal cortex and anterior cingulate cortex. Behav Brain Res 250: 91–101

Bissonette GB, Schoenbaum G, Roesch MR, Powell EM (2015) Interneurons are necessary for coordinated activity during reversal learning in orbitofrontal cortex. Biol Psychiatry 77: 454–64

Blier P, de Montigny C, Chaput Y (1990) A role for the serotonin system in the mechanism of action of antidepressant treatments: preclinical evidence. J Clin Psychiatry 51 Suppl: 14-20; discussion 21

Brigman JL, Daut RA, Wright T, Gunduz-Cinar O, Graybeal C, Davis MI, Jiang Z, Saksida LM, Jinde S, Pease M, Bussey TJ, Lovinger DM, Nakazawa K, Holmes A (2013) GluN2B in corticostriatal circuits governs choice learning and choice shifting. Nat Neurosci 16: 1101–10

Bromberg-Martin ES, Hikosaka O, Nakamura K (2010) Coding of task reward value in the dorsal raphe nucleus. J Neurosci 30: 6262–72

Browne CJ, Abela AR, Chu D, Li Z, Ji X, Lambe EK, Fletcher PJ (2019) Dorsal raphe serotonin neurons inhibit operant responding for reward via inputs to the ventral tegmental area but not the nucleus accumbens: evidence from studies combining optogenetic stimulation and serotonin reuptake inhibition. Neuropsychopharmacology 44: 793–804

Canetta S, Bolkan S, Padilla-Coreano N, Song LJ, Sahn R, Harrison NL, Gordon JA, Brown A, Kellendonk C (2016) Maternal immune activation does not alter the number of perisomatic parvalbumin-positive boutons in the offspring prefrontal cortex. Mol Psychiatry 21: 857

Celada P, Puig MV, Artigas F (2013) Serotonin modulation of cortical neurons and networks. Front Integr Neurosci 7: 25

Chandler DJ, Lamperski CS, Waterhouse BD (2013) Identification and distribution of projections from monoaminergic and cholinergic nuclei to functionally differentiated subregions of prefrontal cortex. Brain Res 1522: 38–58

Cho JR, Treweek JB, Robinson JE, Xiao C, Bremner LR, Greenbaum A, Gradinaru V (2017) Dorsal Raphe Dopamine Neurons Modulate Arousal and Promote Wakefulness by Salient Stimuli. Neuron 94: 1205–1219 e8

Cho KK, Hoch R, Lee AT, Patel T, Rubenstein JL, Sohal VS (2015) Gamma rhythms link prefrontal interneuron dysfunction with cognitive inflexibility in Dlx5/6(+/-) mice. Neuron 85: 1332–43

Chudasama Y, Robbins TW (2003) Dissociable contributions of the orbitofrontal and infralimbic cortex to pavlovian autoshaping and discrimination reversal learning: further evidence for the functional heterogeneity of the rodent frontal cortex. J Neurosci 23: 8771–80

Clarke HF, Dalley JW, Crofts HS, Robbins TW, Roberts AC (2004) Cognitive inflexibility after prefrontal serotonin depletion. Science 304: 878–80

Clarke HF, Walker SC, Crofts HS, Dalley JW, Robbins TW, Roberts AC (2005) Prefrontal serotonin depletion affects reversal learning but not attentional set shifting. J Neurosci 25: 532–8

Cohen JY, Amoroso MW, Uchida N (2015) Serotonergic neurons signal reward and punishment on multiple timescales. Elife 4

Correia PA, Lottem E, Banerjee D, Machado AS, Carey MR, Mainen ZF (2017) Transient inhibition and long-term facilitation of locomotion by phasic optogenetic activation of serotonin neurons. Elife 6

Damasio AR, Tranel D, Damasio H (1990) Individuals with sociopathic behavior caused by frontal damage fail to respond autonomically to social stimuli. Behav Brain Res 41: 81–94

Darrah JM, Stefani MR, Moghaddam B (2008) Interaction of N-methyl-D-aspartate and group 5 metabotropic glutamate receptors on behavioral flexibility using a novel operant set-shift paradigm. Behav Pharmacol 19: 225–34

Davis AK, Barrett FS, Griffiths RR (2020) Psychological flexibility mediates the relations between acute psychedelic effects and subjective decreases in depression and anxiety. J Contextual Behav Sci 15: 39–45

Deacon RM, Penny C, Rawlins JN (2003) Effects of medial prefrontal cortex cytotoxic lesions in mice. Behav Brain Res 139: 139–55

Dennis JP, Vander Wal JS (2010) The Cognitive Flexibility Inventory: Instrument Development and Estimates of Reliability and Validity. Cognitive Therapy and Research 34: 241–253

Dias R, Robbins TW, Roberts AC (1996a) Dissociation in prefrontal cortex of affective and attentional shifts. Nature 380: 69–72

Dias R, Robbins TW, Roberts AC (1996b) Primate analogue of the Wisconsin Card Sorting Test: effects of excitotoxic lesions of the prefrontal cortex in the marmoset. Behav Neurosci 110: 872–86

Dos Santos RG, Bouso JC, Alcazar-Corcoles MA, Hallak JEC (2018) Efficacy, tolerability, and safety of serotonergic psychedelics for the management of mood, anxiety, and substance-use disorders: a systematic review of systematic reviews. Expert Rev Clin Pharmacol 11: 889–902

Elliott MC, Tanaka PM, Schwark RW, Andrade R (2018) Serotonin Differentially Regulates L5 Pyramidal Cell Classes of the Medial Prefrontal Cortex in Rats and Mice. eNeuro 5: ENEURO.0305-17.2018

Fellows LK, Farah MJ (2003) Ventromedial frontal cortex mediates affective shifting in humans: evidence from a reversal learning paradigm. Brain 126: 1830–7

Fernandez SP, Cauli B, Cabezas C, Muzerelle A, Poncer JC, Gaspar P (2016) Multiscale single-cell analysis reveals unique phenotypes of raphe 5-HT neurons projecting to the forebrain. Brain Struct Funct 221: 4007–4025

Fonseca MS, Murakami M, Mainen ZF (2015) Activation of dorsal raphe serotonergic neurons promotes waiting but is not reinforcing. Curr Biol 25: 306–315

Goodwill HL, Manzano-Nieves G, LaChance P, Teramoto S, Lin S, Lopez C, Stevenson RJ, Theyel BB, Moore CI, Connors BW, Bath KG (2018) Early Life Stress Drives Sex-Selective Impairment in Reversal Learning by Affecting Parvalbumin Interneurons in Orbitofrontal Cortex of Mice. Cell Rep 25: 2299–2307 e4

Grossman CD, Bari BA, Cohen JY (2022) Serotonin neurons modulate learning rate through uncertainty. Current Biology 32: 586–599.e7

Heilbronner SR, Rodriguez-Romaguera J, Quirk GJ, Groenewegen HJ, Haber SN (2016) Circuit-Based Corticostriatal Homologies Between Rat and Primate. Biological Psychiatry 80: 509–521

Inaba K, Mizuhiki T, Setogawa T, Toda K, Richmond BJ, Shidara M (2013) Neurons in monkey dorsal raphe nucleus code beginning and progress of step-by-step schedule, reward expectation, and amount of reward outcome in the reward schedule task. J Neurosci 33: 3477–91

Jones JM, Foster W, Twomey CR, Burdge J, Ahmed OM, Pereira TD, Wojick JA, Corder G, Plotkin JB, Abdus-Saboor I (2020) A machine-vision approach for automated pain measurement at millisecond timescales. Elife 9

Kjaerby C, Athilingam J, Robinson SE, Iafrati J, Sohal VS (2016) Serotonin 1B Receptors Regulate Prefrontal Function by Gating Callosal and Hippocampal Inputs. Cell Rep 17: 2882–2890

Klein J, Winter C, Coquery N, Heinz A, Morgenstern R, Kupsch A, Juckel G (2010) Lesion of the medial prefrontal cortex and the subthalamic nucleus selectively affect depression-like behavior in rats. Behav Brain Res 213: 73–81

Kringelbach ML, Rolls ET (2003) Neural correlates of rapid reversal learning in a simple model of human social interaction. Neuroimage 20: 1371–83

Kroes MCW, Dunsmoor JE, Hakimi M, Oosterwaal S, collaboration NP, Meager MR, Phelps EA (2019) Patients with dorsolateral prefrontal cortex lesions are capable of discriminatory threat learning but appear impaired in cognitive regulation of subjective fear. Soc Cogn Affect Neurosci 14: 601-612

Lacroix L, Spinelli S, Heidbreder CA, Feldon J (2000) Differential role of the medial and lateral prefrontal cortices in fear and anxiety. Behav Neurosci 114: 1119–30

Liang B, Zhang L, Barbera G, Fang W, Zhang J, Chen X, Chen R, Li Y, Lin DT (2018) Distinct and Dynamic ON and OFF Neural Ensembles in the Prefrontal Cortex Code Social Exploration. Neuron 100: 700–714 e9

Lisboa SF, Stecchini MF, Correa FM, Guimaraes FS, Resstel LB (2010) Different role of the ventral medial prefrontal cortex on modulation of innate and associative learned fear. Neuroscience 171: 760–8

Liu RJ, Lambe EK, Aghajanian GK (2005) Somatodendritic autoreceptor regulation of serotonergic neurons: dependence on L-tryptophan and tryptophan hydroxylase-activating kinases. Eur J Neurosci 21: 945–58

Lottem E, Banerjee D, Vertechi P, Sarra D, Lohuis MO, Mainen ZF (2018) Activation of serotonin neurons promotes active persistence in a probabilistic foraging task. Nat Commun 9: 1000

Madisen L, Mao T, Koch H, Zhuo JM, Berenyi A, Fujisawa S, Hsu YW, Garcia AJ, 3rd, Gu X, Zanella S, Kidney J, Gu H, Mao Y, Hooks BM, Boyden ES, Buzsaki G, Ramirez JM, Jones AR, Svoboda K, Han X et al. (2012) A toolbox of Cre-dependent optogenetic transgenic mice for light-induced activation and silencing. Nat Neurosci 15: 793-802

Manzano-Nieves G, Liston C (2021) Psychedelics re-engineered for potential use in the clinic. Nature 589: 358–359

Marquardt K, Saha M, Mishina M, Young JW, Brigman JL (2014) Loss of GluN2A-containing NMDA receptors impairs extra-dimensional set-shifting. Genes Brain Behav 13: 611–7

Marton TF, Seifikar H, Luongo FJ, Lee AT, Sohal VS (2018) Roles of Prefrontal Cortex and Mediodorsal Thalamus in Task Engagement and Behavioral Flexibility. J Neurosci 38: 2569–2578

Matias S, Lottem E, Dugue GP, Mainen ZF (2017) Activity patterns of serotonin neurons underlying cognitive flexibility. Elife 6

Miyazaki K, Miyazaki KW, Sivori G, Yamanaka A, Tanaka KF, Doya K (2020) Serotonergic projections to the orbitofrontal and medial prefrontal cortices differentially modulate waiting for future rewards. Sci Adv 6

Miyazaki K, Miyazaki KW, Yamanaka A, Tokuda T, Tanaka KF, Doya K (2018) Reward probability and timing uncertainty alter the effect of dorsal raphe serotonin neurons on patience. Nat Commun 9: 2048

Miyazaki KW, Miyazaki K, Tanaka KF, Yamanaka A, Takahashi A, Tabuchi S, Doya K (2014) Optogenetic activation of dorsal raphe serotonin neurons enhances patience for future rewards. Curr Biol 24: 2033–40

Murphy-Beiner A, Soar K (2020) Ayahuasca’s ‘afterglow’: improved mindfulness and cognitive flexibility in ayahuasca drinkers. Psychopharmacology (Berl) 237: 1161–1169

Muzerelle A, Scotto-Lomassese S, Bernard JF, Soiza-Reilly M, Gaspar P (2016) Conditional anterograde tracing reveals distinct targeting of individual serotonin cell groups (B5-B9) to the forebrain and brainstem. Brain Struct Funct 221: 535–61

Nakamura K, Matsumoto M, Hikosaka O (2008) Reward-dependent modulation of neuronal activity in the primate dorsal raphe nucleus. J Neurosci 28: 5331–43

Nutt DJ (2005) Overview of diagnosis and drug treatments of anxiety disorders. CNS Spectr 10: 49–56

O’Doherty J, Critchley H, Deichmann R, Dolan RJ (2003) Dissociating valence of outcome from behavioral control in human orbital and ventral prefrontal cortices. J Neurosci 23: 7931–9

Ohmura Y, Tanaka KF, Tsunematsu T, Yamanaka A, Yoshioka M (2014) Optogenetic activation of serotonergic neurons enhances anxiety-like behaviour in mice. Int J Neuropsychopharmacol 17: 1777–83

Prouty EW, Chandler DJ, Waterhouse BD (2017) Neurochemical differences between target-specific populations of rat dorsal raphe projection neurons. Brain Res 1675: 28–40

Ragozzino ME (2007) The contribution of the medial prefrontal cortex, orbitofrontal cortex, and dorsomedial striatum to behavioral flexibility. Ann N Y Acad Sci 1121: 355–75

Ranade SP, Mainen ZF (2009) Transient firing of dorsal raphe neurons encodes diverse and specific sensory, motor, and reward events. J Neurophysiol 102: 3026–37

Rebello TJ, Yu Q, Goodfellow NM, Caffrey Cagliostro MK, Teissier A, Morelli E, Demireva EY, Chemiakine A, Rosoklija GB, Dwork AJ, Lambe EK, Gingrich JA, Ansorge MS (2014) Postnatal day 2 to 11 constitutes a 5-HT-sensitive period impacting adult mPFC function. J Neurosci 34: 12379–93

Rex A, Fink H (1998) Effects of cholecystokinin-receptor agonists on cortical 5-HT release in guinea pigs on the X-maze. Peptides 19: 519–26

Rogers RD, Andrews TC, Grasby PM, Brooks DJ, Robbins TW (2000) Contrasting cortical and subcortical activations produced by attentional-set shifting and reversal learning in humans. J Cogn Neurosci 12: 142–62

Rojas PS, Fiedler JL (2016) What Do We Really Know About 5-HT1A Receptor Signaling in Neuronal Cells? Front Cell Neurosci 10: 272

Roy JE, Riesenhuber M, Poggio T, Miller EK (2010) Prefrontal cortex activity during flexible categorization. J Neurosci 30: 8519–28

Rygula R, Walker SC, Clarke HF, Robbins TW, Roberts AC (2010) Differential contributions of the primate ventrolateral prefrontal and orbitofrontal cortex to serial reversal learning. J Neurosci 30: 14552–9

Sargin D, Chottekalapanda RU, Perit KE, Yao V, Chu D, Sparks DW, Kalik S, Power SK, Troyanskaya OG, Schmidt EF, Greengard P, Lambe EK (2020) Mapping the physiological and molecular markers of stress and SSRI antidepressant treatment in S100a10 corticostriatal neurons. Mol Psychiatry 25: 1112–1129

Sargin D, Jeoung HS, Goodfellow NM, Lambe EK (2019) Serotonin Regulation of the Prefrontal Cortex: Cognitive Relevance and the Impact of Developmental Perturbation. ACS Chem Neurosci 10: 3078–3093

Scott MM, Wylie CJ, Lerch JK, Murphy R, Lobur K, Herlitze S, Jiang W, Conlon RA, Strowbridge BW, Deneris ES (2005) A genetic approach to access serotonin neurons for in vivo and in vitro studies. Proc Natl Acad Sci U S A 102: 16472–7

Seo C, Ito B, Krupa NA, Jin M, Guru A, Wang E, Shen CX, Boada C, Kullakanda DS, Ho Y-Y, Sleezer BJ, Warden MR (2019) Intense threat switches dorsal raphe serotonin neurons to a paradoxical operational mode. Science, 363(6426), 538–542.

Shah AA, Treit D (2003) Excitotoxic lesions of the medial prefrontal cortex attenuate fear responses in the elevated-plus maze, social interaction and shock probe burying tests. Brain Res 969: 183–94

Shallice T, Stuss DT, Picton TW, Alexander MP, Gillingham S (2007) Multiple effects of prefrontal lesions on task-switching. Front Hum Neurosci 1: 2

Sparks DW, Tian MK, Sargin D, Venkatesan S, Intson K, Lambe EK (2017) Opposing Cholinergic and Serotonergic Modulation of Layer 6 in Prefrontal Cortex. Front Neural Circuits 11: 107

Spellman T, Svei M, Kaminsky J, Manzano-Nieves G, Liston C (2021) Prefrontal deep projection neurons enable cognitive flexibility via persistent feedback monitoring. Cell 184: 2750–2766 e17

Stuss DT (2011) Functions of the frontal lobes: relation to executive functions. J Int Neuropsychol Soc 17: 759–65

Sullivan RM, Gratton A (2002) Behavioral effects of excitotoxic lesions of ventral medial prefrontal cortex in the rat are hemisphere-dependent. Brain Res 927: 69–79

Szczepanski SM, Knight RT (2014) Insights into human behavior from lesions to the prefrontal cortex. Neuron 83: 1002–18

Teissier A, Chemiakine A, Inbar B, Bagchi S, Ray RS, Palmiter RD, Dymecki SM, Moore H, Ansorge MS (2015) Activity of Raphe Serotonergic Neurons Controls Emotional Behaviors. Cell Rep 13: 1965–76

Teixeira CM, Rosen ZB, Suri D, Sun Q, Hersh M, Sargin D, Dincheva I, Morgan AA, Spivack S, Krok AC, Hirschfeld-Stoler T, Lambe EK, Siegelbaum SA, Ansorge MS (2018) Hippocampal 5-HT Input Regulates Memory Formation and Schaffer Collateral Excitation. Neuron 98: 992–1004 e4

Tian MK, Schmidt EF, Lambe EK (2016) Serotonergic Suppression of Mouse Prefrontal Circuits Implicated in Task Attention. eNeuro 3

Torrado Pacheco A, Olson RJ, Garza G, Moghaddam B (2023) Acute psilocybin enhances cognitive flexibility in rats. Neuropsychopharmacology

Troudet R, Detrait E, Hanon E, Lamberty Y (2016) Optimization and pharmacological validation of a set-shifting procedure for assessing executive function in rats. J Neurosci Methods 268: 182–8

Winkelman M (2014) Psychedelics as medicines for substance abuse rehabilitation: evaluating treatments with LSD, Peyote, Ibogaine and Ayahuasca. Curr Drug Abuse Rev 7: 101–16

Winstanley CA, Chudasama Y, Dalley JW, Theobald DE, Glennon JC, Robbins TW (2003) Intra-prefrontal 8-OH-DPAT and M100907 improve visuospatial attention and decrease impulsivity on the five-choice serial reaction time task in rats. Psychopharmacology (Berl) 167: 304–14

Yamashita PSM, Rosa DS, Lowry CA, Zangrossi H (2018) Serotonin actions within the prelimbic cortex induce anxiolysis mediated by serotonin 1a receptors. Journal of Psychopharmacology 33: 3–11

Yin L, Rasch MJ, He Q, Wu S, Dou F, Shu Y (2017) Selective Modulation of Axonal Sodium Channel Subtypes by 5-HT1A Receptor in Cortical Pyramidal Neuron. Cereb Cortex 27: 509–521

Zhong P, Yan Z (2011) Differential regulation of the excitability of prefrontal cortical fast-spiking interneurons and pyramidal neurons by serotonin and fluoxetine. PLoS One 6: e16970

