## supplementary figures and table for "Medial Prefrontal Cortex Serotonin Input Regulates Cognitive Flexibility in Mice"

### Supplementary Information

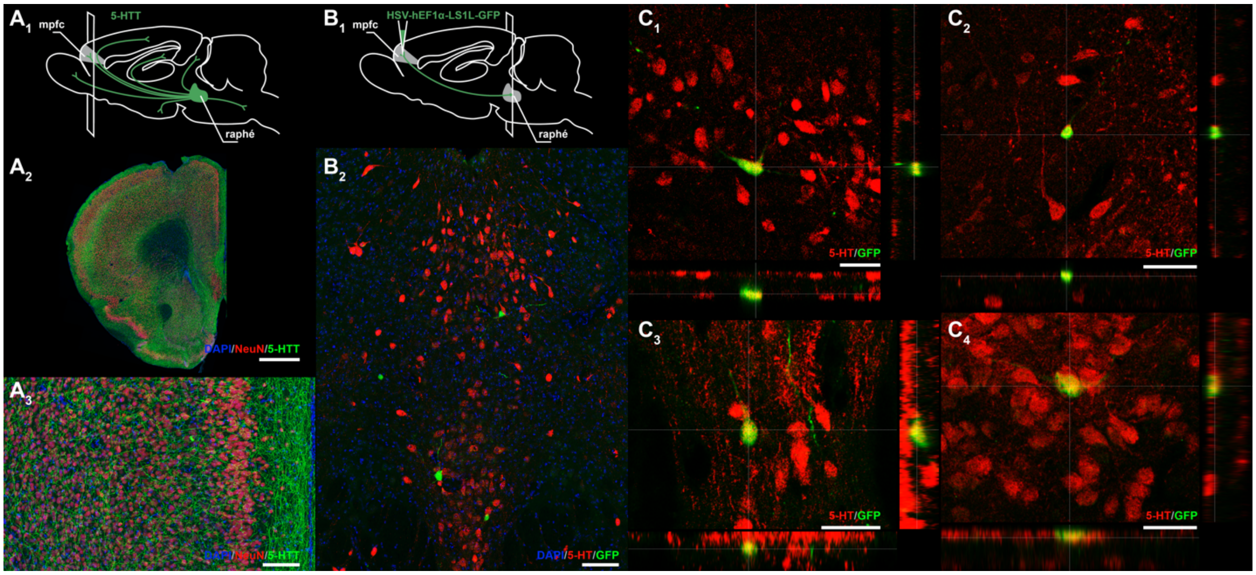


**Supplementary Figure 1. DR serotonergic neurons project to the PFC.** The mPFC receives dense serotonergic innervation, as revealed by the double-staining of the neuronal marker NeuN (red) and the serotonin transporter (5-HTT, green) (**A1**, **A2**). Viral retrograde tracing using HSV-hEF1α-LS1L-GFP into the mPFC of SERT-cre mice (**B1**) reveals retrogradely labeled serotonergic neurons in the DR (**B2**). GFP-positive neurons (green) co-label with the serotonergic marker 5-HT (red). Scale bars 1 mm (**A2**), 100 um (**A3, B2**) and 50 um (**C1-4**).

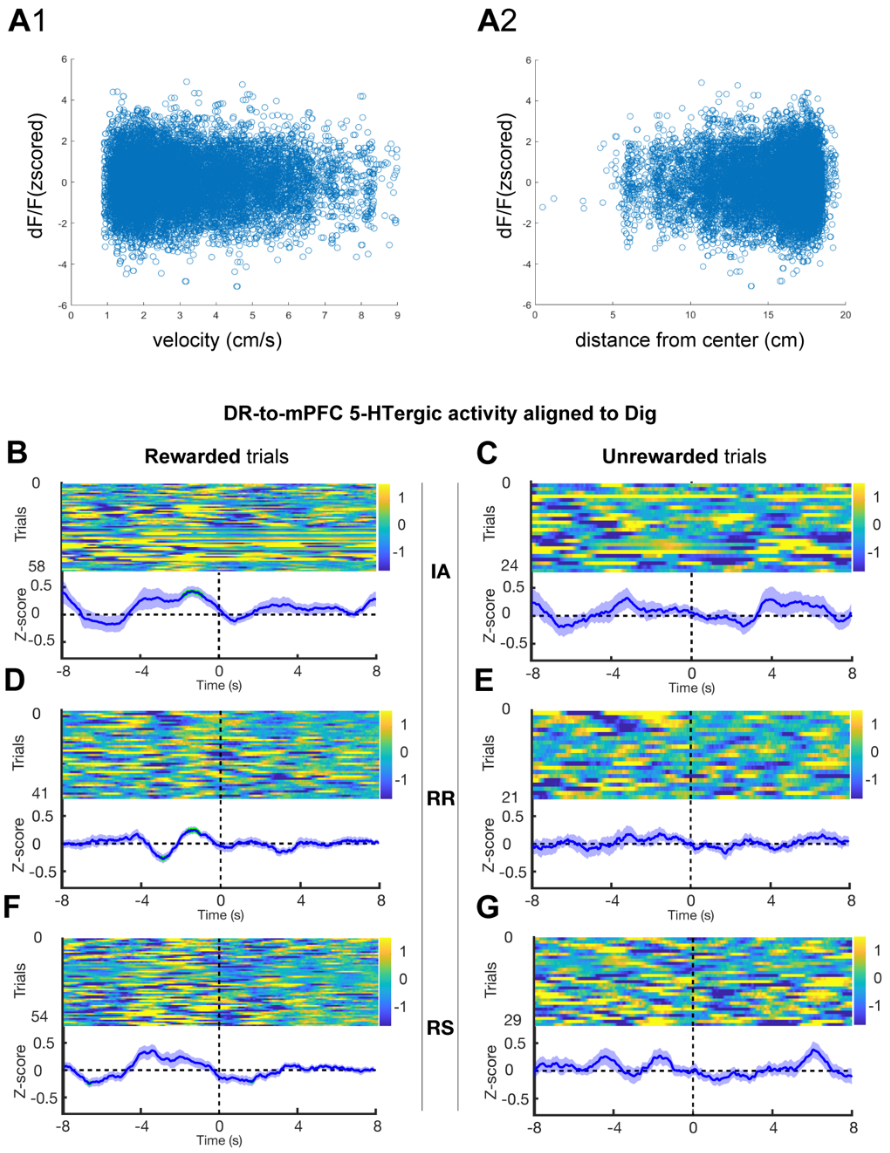


**Supplementary Figure 2. Activity signatures of mPFC-projecting DR serotonergic neurons in the open-field test and two-choice digging test.** Exemplary individual correlations between the activity of DR-to-mPFC serotonergic neurons, using fiber photometry, and velocity (**A**1) or distance in the center (**A**2) in the open-field test. The activity of mPFC-projecting DR serotonergic neurons in the two-choice digging tasks during IA, RR, and RS aligned to the dig initiation (choice) in successful and unsuccessful trials (**B-G**), n = 4-6/group.

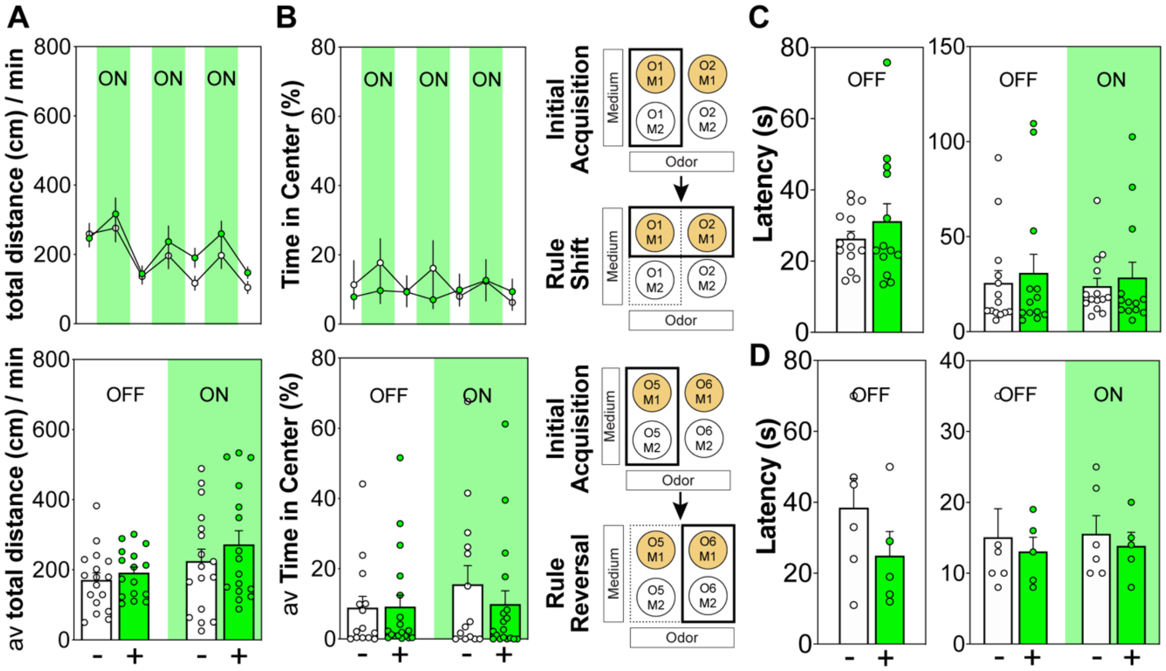


**Supplementary Figure 3. Optogenetic inhibition of serotonergic terminals in the PL produces no changes in time in center and total distance in the open-field test or time to decision in the two-choice digging test.** Optogenetic inhibition of serotonergic terminals in the PL did not impact time in center (**A**) and total distance traveled (**B**) in the open-field test. Similarly, no changes in the latency to dig initiation (decision) were observed as a consequence of opto-inhibition during the extradimensional rule shift (**C**) and intradimensional rule shift (**D**) task of the two-choice digging test. N = 13-18/group.

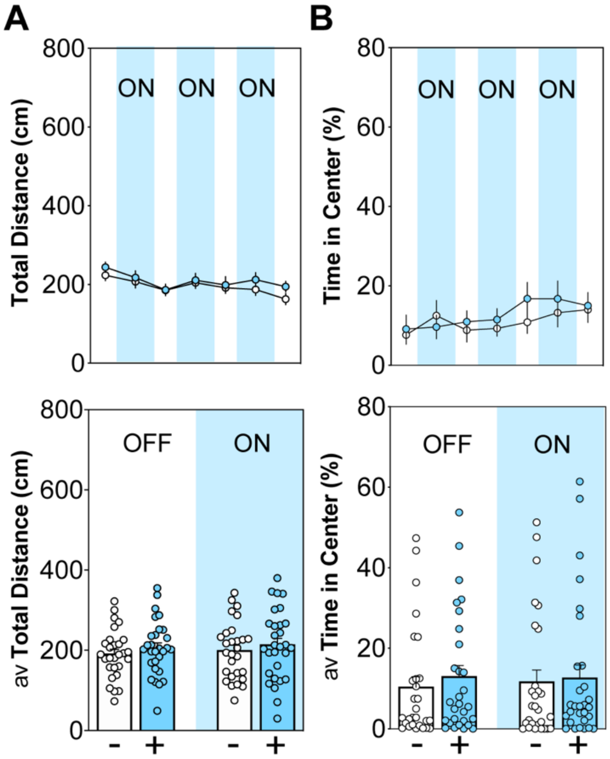


**Supplementary Figure 4. Optogenetic activation of serotonergic terminals in the PL does not impact on anxiety-like behavior and locomotor activity in the open-field test.** Opto-stimulation of PL serotonergic terminals produced no changes in total distance (**A**) and time in center (**B**) in the open-field test. N = 12-21/group.

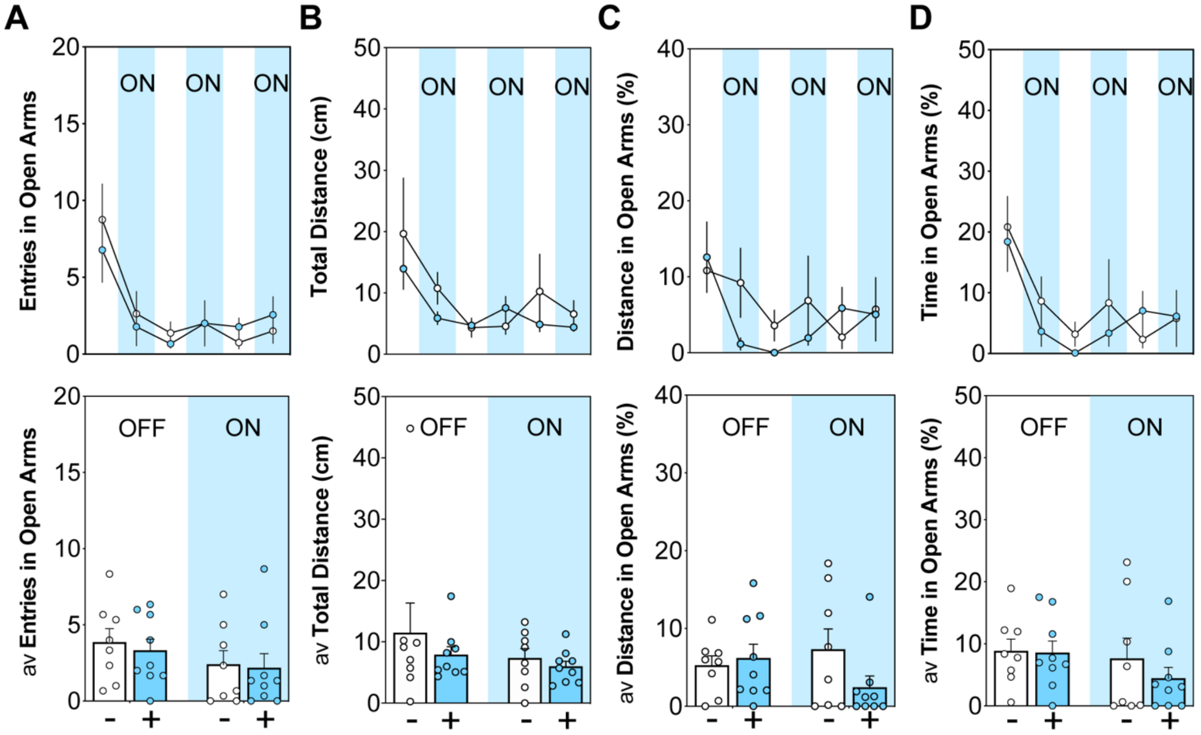


**Supplementary Figure 5. Optogenetic activation of serotonergic terminals in the PL does not change locomotor activity and anxiety-like behavior in the elevated-plus maze.** Opto-stimulation of PL serotonergic terminals does not change entries into open arms (**A**), total distance traveled (**B**), distance traveled in open arms (**C**), or time spent in the open arms (**D**) of elevated-plus maze test. N = 8-9/group.

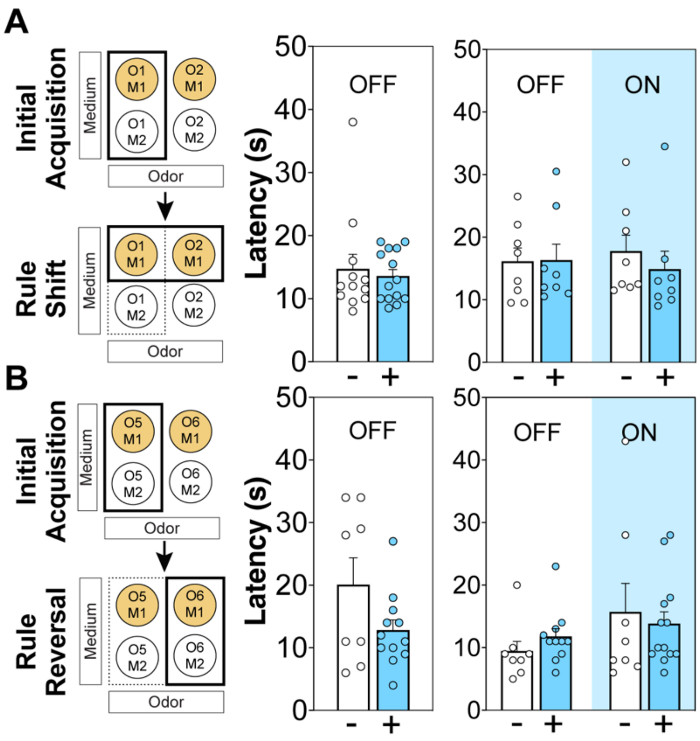


**Supplementary Figure 6. Optogenetic activation of serotonergic terminals in the PL does not impact latency to decision in cognitive flexibility tasks.** No changes in the latency to dig initiation (decision) were observed as a consequence of opto-stimulation during the extradimensional rule shift (**A**) and intradimensional rule shift (**B**) task of the two-choice digging test. N = 12-21/group.

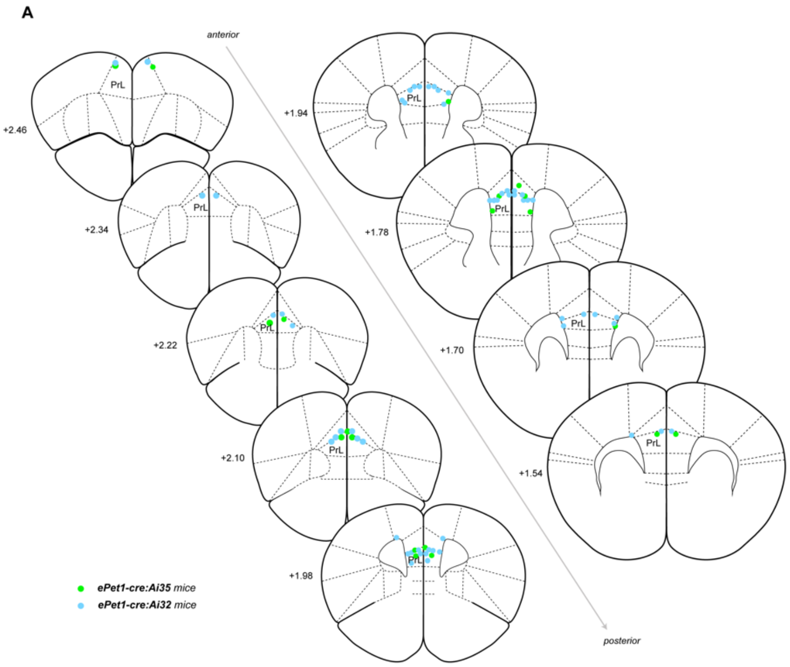


**Supplementary Figure 7: Fiber placements from optogenetic experiments.**

| | | | Dimensions | | Combinations | | Stimulation Group | |
|---|---|---|---|---|---|---|---|---|
| Day | Order | Task | **Relevant** | Irrelevant | **Correct** | Incorrect | 1 | 2 |
| 1 | A | IA (1) | **Odor** | Texture | **Cinnamon**/Paper | Paprika/ Paper | OFF | OFF |
| | | | | | **Cinnamon**/Corn | Paprika/ Corn | | |
| | B | RS (1) | Texture | Odor | Cinnamon/**Paper** | Cinnamon/Corn | **ON** | OFF |
| | | | | | Paprika/ **Paper** | Paprika/ Corn | | |
| 2 | A | IA (2) | Texture | Odor | Cinnamon/**Paper** | Cinnamon/Corn | OFF | OFF |
| | | | | | Paprika/ **Paper** | Paprika/ Corn | | |
| | B | RS (2) | **Odor** | Texture | **Paprika**/ Paper | Cinnamon/ Paper | OFF | **ON** |
| | | | | | **Paprika**/ Corn | Cinnamon/Corn | | |
| 3 | A | IA (3) | **Odor** | Texture | **Paprika**/ Paper | Cinnamon/ Paper | OFF | OFF |
| | | | | | **Paprika**/ Corn | Cinnamon/Corn | | |
| | B | RS (3) | Texture | Odor | Cinnamon/**Corn** | Cinnamon/Paper | OFF | **ON** |
| | | | | | Paprika/ **Corn** | Paprika/ Paper | | |
| 4 | A | IA (4) | Texture | Odor | Cinnamon/**Corn** | Cinnamon/Paper | OFF | OFF |
| | | | | | Paprika/ **Corn** | Paprika/ Paper | | |
| | B | RS (4) | **Odor** | Texture | **Cinnamon**/Paper | Paprika/ Paper | **ON** | OFF |
| | | | | | **Cinnamon**/Corn | Paprika/ Corn | | |
| 5 | A | IA (5) | **Odor** | Texture | **Cinnamon**/Paper | Paprika/ Paper | OFF | OFF |
| | | | | | **Cinnamon**/Corn | Paprika/ Corn | | |
| | B | RR (1) | **Odor** | Texture | **Paprika**/ Paper | Cinnamon/ Paper | **ON** | OFF |
| | | | | | **Paprika**/ Corn | Cinnamon/Corn | | |
| 6 | A | IA (6) | **Odor** | Texture | **Paprika**/ Paper | Cinnamon/ Paper | OFF | OFF |
| | | | | | **Paprika**/ Corn | Cinnamon/Corn | | |
| | B | RR (2) | **Odor** | Texture | **Cinnamon**/Paper | Paprika/ Paper | OFF | **ON** |
| | | | | | **Cinnamon**/Corn | Paprika/ Corn | | |

**Supplementary Table 1: Sequences of odor and texture associations.**
